# Disease-specific tangle immunophenotypes distinguish hippocampal vulnerability in Alzheimer’s disease and Parkinson’s disease dementia

**DOI:** 10.64898/2026.08.13.744594

**Authors:** Sophie Schreiner, Mónica Miranda de la Maza, Gaël Paul Hammer, Félicia Jeannelle, Morgane Darricau, Dominique Mirault, Naguib Mechawar, Netherlands Brain Bank, Michel Mittelbronn, David S. Bouvier

## Abstract

INTRODUCTION

Tau pathology typically occurs in Alzheimeŕs disease (AD), however is also frequently present in Parkinsońs disease dementia (PDD) and Dementia with Lewy Bodies (DLB), yet its disease-specific signature is unclear.

**METHODS:** Five tau, amyloid-β, α-synuclein and neuronal markers were analysed across hippocampal subfields in non-demented controls (CTLs), AD, PDD and DLB using multiplex immunohistochemistry, single-tangle classification and confocal imaging.

**RESULTS:** AT8, pTau217, and GT38 were predominatly detected in AD, while pS422 was enriched in PDD and pS396 showed a region- and disease-specific pattern. DLB resembled AD in subregional tau distribution. Tau marker correlation were different comparing AD, PDD and CTL. Single-tangle analyses revealed disease-specific immunophenotypes but conserved mature intra-tangle epitope organisation. Distinct tau signatures were associated with inhibitory interneuron vulnerability, while regional tau co-occurrence with amyloid-β and α-synuclein remained conserved.

**DISCUSSION:** Disease-specific tau signatures vary across hippocampal subregions and neuronal populations, implicating the contribution of regional and cell-specific factors beyond pathology burden.

## 1 Background

Memory decline is a debilitating and defining clinical feature of neurodegenerative diseases (NDDs) [1]. Despite differences in cause, genetics and primary pathology, Alzheimeŕs disease (AD), Parkinsońs disease dementia (PDD) and Dementia with Lewy bodies (DLB) converge on progressive memory and cognitive impairment [2,3]. The hippocampus is central to memory encoding and consolidation, and hippocampal neuronal integrity is linked to cognitive decline severity [4]. The extent of hippocampal deterioration across these disorders remains debated, particularly given reports of relative hippocampal preservation in Lewy body disorders [5].

Tau pathology, a hallmark of AD, is a strongly associated with hippocampal neurodegeneration [6]. It is also present in other NDDs, including DLB, although its contribution to PDD neurodegeneration remains unclear [7]. Under physiological conditions, tau is a neuronal microtubule-associated protein that stabilises axonal microtubules [8]. In disease, abnormally hyper-phosphorylated tau dissociates from microtubules and aggregates into paired helical filaments that condense into neurofibrillary tangles (NFTs) [9]. NFTs maturation, from early intraneuronal pretangles, through mature tangles to extracellular ghost tangles, involves sequential acquisition of specific phosphorylation epitopes and conformational changes [10]. Tau antibodies show differential immunoreactivity across NFT maturation stages. Early pretangle stages are characterised by phosphorylation at sites including pS422, whereas mature tangles accumulate AT8 (pS202/pT205) and pS396 epitopes. Late conformational changes are captured by conformation-sensitive antibodies as GT38 [10,11]. The stereotypical progression of NFT pathology through defined brain regions, from the transentorhinal cortex through hippocampal subfields to neocortical areas, was first described by Braak and Braak, based on AT8 immunoreactivity and underlies the Braak staging system [12]. Recent biochemical work has further refined this framework by distinguishing phosphorylation states associated with soluble, pre- fibrillar tau assemblies from the mature, fibrillar NFT [13]. Whether pathological tau states define spatial microenvironments or cellular vulnerability across tauopathies remains largely unexplored.

Although AD and PDD differ in their primary pathological context, both exhibit tau pathology, with PDD showing tau accumulation alongside synuclein burden [3,14,15]. Whether tau pathology in PDD shares the molecular and spatial characteristics of an early AD tau pathology, or represents a distinct pathological entity shaped by the α- synuclein disease environment, remains unknown. Tau accumulation in dopaminergic neurons independent of nigrostriatal pathology suggests that tau alterations are not solely secondary to α-synuclein pathology in Parkinsońs disease (PD) [16]. DLB provides an important comparative model, given its clinical and pathological overlap with both PD and AD [15], and highlights the need to define whether tau pathology exhibits disease-specific features across disorders [17–19].

The hippocampus is of particular interest in this context, as it represents a major substrate of the episodic memory decline that characterise the transition from PD to PDD, and accumulates both tau and α-synuclein pathology during disease progression [20]. However, the extent to which hippocampal tau in PDD recapitulates the phosphorylation trajectory, tangle composition, and neuronal vulnerability profile observed in AD remains poorly characterised. Key unresolved questions include PDD tau maturation, α-synuclein effects on tau organisation and whether both pathologies converge within shared neuronal populations or define distinct vulnerability patterns. Addressing these questions will clarify the shared and distinct neuropathological mechanisms underlying cognitive decline in PDD and AD, with implications for tau-directed therapeutic strategies across dementia syndromes.

To address these questions, we performed a quantitative characterisation of the hippocampal tau pathology, neuronal integrity and co-pathology in *post-mortem* cohorts comprising age-matched non-demented control (CTL), AD, PDD and DLB cases. Using tau-epitope panel spanning stages of NFT maturation (pS422, AT8, pS396, pTau217 and GT38) together with neuronal, synaptic, amyloid-β and α- synuclein markers, we assessed whether tau burden, tau composition, neuronal vulnerability and pathological interactions differ across disorders. Regional quantification across hippocampal subfields and the parahippocampal cortex (PHC) was integrated with correlation analysis, single-tangle multiplex co-staining, and high-resolution confocal imaging to define the molecular architecture of tau pathology across spatial scales [21].

## 2 Methods

### 2.1 Study design and workflow

Hippocampal tissue was processed from formalin-fixed paraffin-embedded (FFPE) sections through single-marker immunohistochemistry combined with digital pathology quantification, followed by chromogenic multiplex immunohistochemistry and multiplex immunofluorescence for further visualisation of marker co-localisation (**Fig. 1A**) [21]. Five tau markers were selected to capture distinct stages and conformations along the tangle maturation sequence: pS422 (early phosphorylation), AT8 (broad hyperphosphorylation), pS396 (mature phosphorylation), pTau217 (biomarker-relevant late epitope), and GT38 (conformational change) [10,12,22,23]. These were assessed alongside five neuronal and synaptic markers (NeuN, VGAT, VGLUT1, SST, PVALB) and co-pathology markers for amyloid-β (4G8) and α-synuclein (pSyn81A). Representative staining and digital pathology threshold for each marker are shown in **Fig. 1B**.

**Figure 1.**
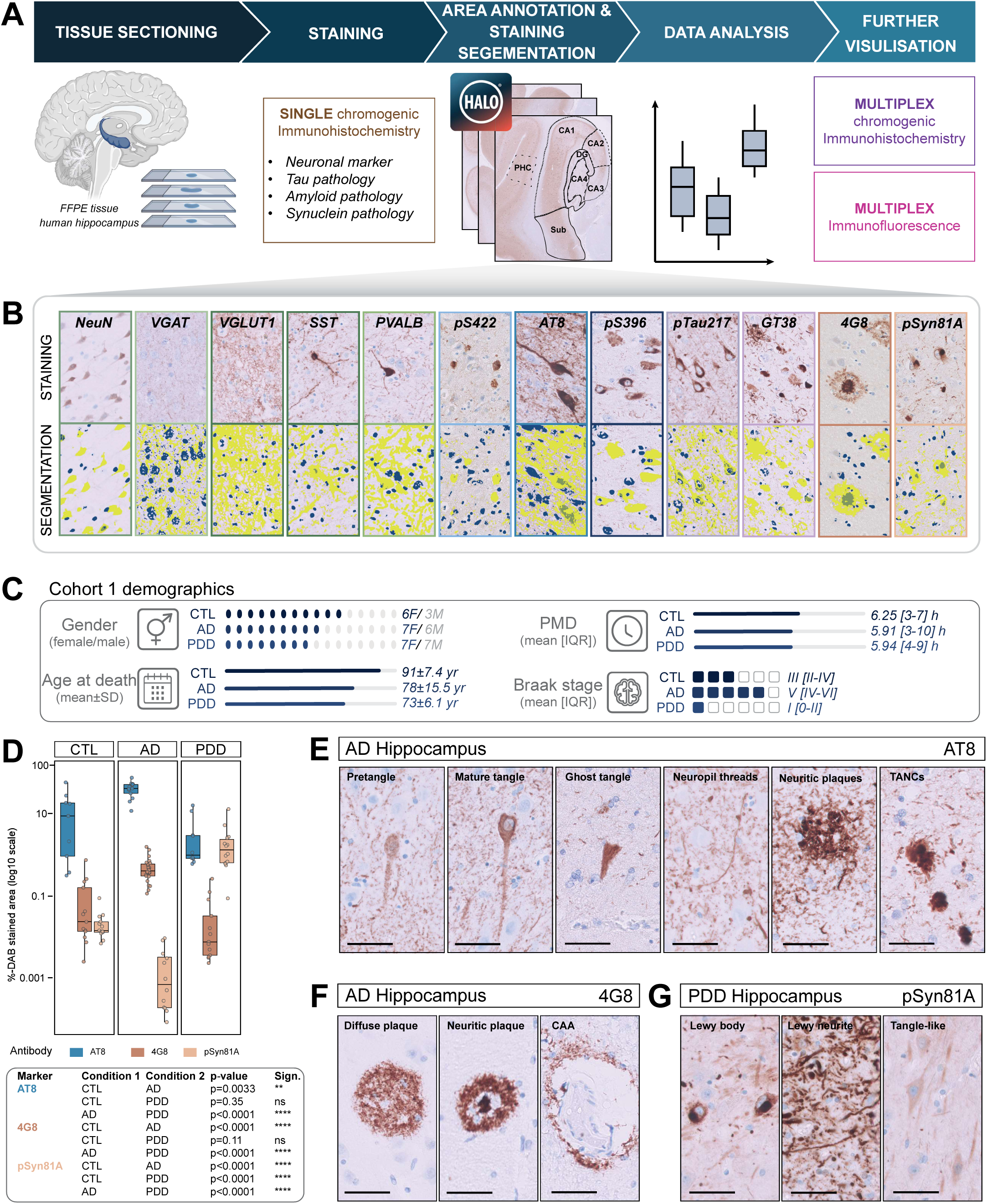
Cohort 1 characterisation and disease-specific co-pathology of CTL, AD, and PDD hippocampal samples. **A** Workflow schematic of tissue processing, staining, and digital pathology analysis. **B** Representative illustration of DAB stain colour thresholding, optimised individually per marker to account for variability in staining intensity and distribution. Underlying segmentation parameters are listed in Supplementary Table 3. **C** Demographic and neuropathological characteristics of cohort 1 (CTL n = 9, AD n=13, PDD n=14) including gender distribution, age at death, *post-mortem* delay (PMD), and Braak NFT stage. **D** Regional AT8, 4G8, and pSyn81A immunoreactivity (%-DAB stained area) at the whole-hippocampus level in CTL, AD, and PDD. **E** Representative images illustrate the staining pattern of AT8 in cIHC including pretangles, mature tangles, ghost tangles, neuropil threads, neuritic plaques and tangle associated neuritic clusters (TANCs) in AD (case#20) **F** Representative 4G8 cIHC illustrating diffuse plaques, neuritic plaques, and cerebral amyloid angiopathy (CAA) in AD (case#32). **G** Representative pSyn81A cIHC illustrating Lewy body, Lewy neurite, and tangle-like α- synuclein morphology in PDD (case#57). Scale bars: 50 µm.

### 2.2 *Post-mortem* human brain samples

This study was conducted using human *post-mortem* formalin-fixed paraffin-embedded (FFPE) hippocampal samples obtained from the Douglas-Bell Canada Brain Bank (Douglas Mental Health University Institute, Montreal, QC, Canada), Netherlands Brain Bank (NBB, Amsterdam, Netherlands) and the GIE-Neuro-CEB biobank (Groupe Hospitalier Pitié-Salpêtrière, Paris, France). Neuropathological characterisation was performed by expert neuropathologists at the respective brain banks, applying established diagnostic frameworks, including Braak staging [12] and the ABC score [24]. The use of these samples was approved by the corresponding institutional ethics committees, as well as by the University of Luxembourg (ERP 16- 037 and 21-009). For this study, FFPE hippocampal tissue was analysed from age-matched non-demented controls (CTL; cohort 1 n=9; cohort 2 n=8), Alzheimer’s disease (AD; cohort 1 n=12; cohort 2 n=11), Parkinson’s disease dementia (PDD; cohort 1 n=14), Dementia with Lewy bodies (DLB; cohort 2 n=10), Frontotemporal Lobar degeneration with Pick-type tau pathology (3R-tau FTLD; n=3) and Progressive supranuclear palsy (PSP; n=3). Selection criteria included age at death, gender, post-mortem delay (PMD), and neuropathological stage (**Supplementary Table 1**).

### 2.3 Single chromogenic immunohistochemistry (scIHC)

FFPE hippocampal tissues were sectioned at 5 µm thickness using a microtome and mounted on glass slides following an incubation at 60°C for 1 hour. Depending on the automated platform used for scIHC, tissue sections were distributed onto separate slides for processing either on the Dako Omnis Immunostainer (Agilent Technologies) using Dako FLEX IHC-coated slides (Cat# K8020, Agilent) or the Ventana Discovery Ultra (Roche Diagnostics) using hydrophilic adhesion slides (Matsunami TOMO®, Cat# TOM-1190).

For each antibody, an optimized staining protocol was applied according to the specifications of the respective platform. In brief, sections underwent automated deparaffinisation and rehydration, followed by heat-induced epitope retrieval (HIER) with buffer conditions tailored to the target antigen. Endogenous peroxidase activity was blocked using hydrogen peroxide. Primary antibodies were applied at validated working dilutions, with incubation times and temperatures adjusted according to the antibody-specific protocol. Detection was performed using HRP-conjugated secondary antibodies and visualized with 3,3′-diaminobenzidine (DAB) as the chromogenic substrate.

For the IHC performed on the Dako Omnis Immunostainer (Agilent Technologies), sections were subjected to HIER for 30 min at 97 °C using either high- or low-pH buffer solutions (EnVision™ FLEX Target Retrieval Solution, high pH: Cat# K8004; low pH: Cat# K8005), depending on the antibody (details provided in **Supplementary Table 2**). Primary antibodies against 4G8, pSyn81A, AT8, pS422, pS396, and GT38 were diluted in EnVision™ FLEX Antibody Diluent (Cat# K8006) at the concentrations listed in **Supplementary Table 2**, and incubated with the tissue sections for 1 h at room temperature (RT). Detection was carried out using the EnVision FLEX Detection Kit (Cat# K8000) according to the manufacturer’s guidelines and protocol.

For scIHC staining on the Ventana Discovery Ultra platform (Roche Diagnostics), paraffin sections were initially deparaffinized at 69 °C in EZ Prep solution (Cat# 950- 102), applied in three consecutive 8-min cycles. Antigen retrieval was subsequently performed at 95 °C for 40 min using Cell Conditioning 1 buffer (CC1, Cat# 950-224). To suppress endogenous peroxidase activity and reduce nonspecific binding, slides were treated with ChromoMap (CM) inhibitor for 8 min at 37 °C. Primary antibodies targeting pTau217, NeuN, SST, PVALB, VGAT and VGLUT1, were applied manually after dilution in EnVision™ FLEX buffer (details provided in **Supplementary Table 2**). Tissue sections were incubated with the primary antibodies for 60 min. Detection was performed using the OmniMap anti-Rabbit (Cat# 760-4311) or OmniMap anti-Mouse (Cat# 760-4310) secondary reagents with a 16-min incubation. The chromogenic reaction was achieved sequentially with H₂O₂ CM (4 min), DAB CM (8 min), and Copper CM (4 min), all components of the Discovery CM DAB kit (Cat# 760-159). Finally, all slides were counterstained with haematoxylin, dehydrated through graded ethanol and xylene, and mounted with coverslips. Each antibody staining included prior positive and negative controls processed under identical conditions to ensure staining quality and specificity.

### 2.4 Quantification method and statistical analyses

Manual tangle quantification was done using QuPath [25] and manual annotations. The tissue regions of interest were manually annotated based on *Atlas of the human brain* [26]. Each tangle of each region was then manually classified into one of seven possible immunophenotypes. Statistical analyses were performed using R Studio software (R Foundation for Statistical Computing, Vienna, Austria). For each patient, tangle counts were grouped by slide, region, and phenotype, and converted into within-region percentages. These values were then completed across all phenotype/region/patient combinations and averaged to obtain the mean, standard deviation, and patient count for each phenotype within each region.

Quantitative analysis of IHC staining was performed using HALO image analysis software (v4.0.5107.445, Indica Labs, Albuquerque, NM, USA). The HALO® Area Quantification module was applied to determine the extent of chromogen-positive labelling. General analysis parameters were kept constant across all stainings, including HTX stain colour (0.097; 0.097; 0.066), HTX Min OD (0.078; 0.1496; 0.3165), background stain colour (0.058; 0.066; 0.054), and minimum tissue OD (0.037). For each antibody, thresholds for DAB stain colour and DAB minimum OD were optimized individually to account for variability in DAB staining intensity and distribution (**Fig. 1B**). This optimization was performed using reference slides representing low and high levels of immunoreactivity, and the final settings are summarized in **Supplementary Table 3**. For every case and antibody, tissue regions of interest were manually annotated based on *Atlas of the human brain* [26] to delineate the dentate gyrus (DG), *cornu ammonis* subfields (CA1–CA4), subiculum, and parahippocampal cortex (PHC).

Following annotation, HALO automatically quantified the stained and unstained tissue within each region. Although the raw output generated by HALO includes multiple measurements, only the total annotated tissue area (“Tissue Area Analyzed (µm²)”) and the DAB-positive area (“DAB Area (µm²)“) were retained for further analysis. Statistical analyses were performed using R Studio software (R Foundation for Statistical Computing, Vienna, Austria). For each region of interest and antibody, the stained area percentage was calculated as the ratio of the DAB-positive area to the total annotated tissue area. Data were visualized as boxplots to illustrate group distributions. Group comparisons were carried out using the Wilcoxon rank-sum test, with statistical significance set at p< 0.05. Levels of significance are indicated as follows: p< 0.05 (*), p< 0.005 (**), and p< 0.0005 (***).

To assess the relative balance of GABAergic and glutamatergic markers, measurements of VGAT and VGLUT1 were analysed across the anatomical regions included in the dataset, with the PHC region excluded from the analysis. VGAT and VGLUT1 DAB-positive tissue measurements were converted to numeric values and paired within each sample to calculate the VGAT/VGLUT1 ratio. Ratios were averaged within each condition and anatomical region and subsequently log2-transformed to represent the relative balance between the two markers. Mean log2-transformed ratios were visualised using heatmaps across conditions and anatomical regions using R Studio software (R Foundation for Statistical Computing, Vienna, Austria) and Adobe Illustrator.

### 2.5 Multiplex chromogenic immunohistochemistry (mcIHC)

Multiplex cIHC was performed to enable the sequential chromogenic detection of multiple targets within the same tissue section using the Ventana Discovery Ultra platform (Roche Diagnostics), as previously described. All multiplex protocols (two-, three-, and four-plex) shared the same initial steps of deparaffinisation, heat-induced epitope retrieval (HIER) using Cell Conditioning 1 (CC1), and endogenous enzyme blocking with Discovery Inhibitor CM, as described for single-plex staining. For all multiplex panels, one drop of Discovery Inhibitor (Cat# 760–4840) was applied for 8 min following deparaffinisation and HIER. After each staining cycle, heat-mediated antibody denaturation was performed using Cell Conditioning 2 (CC2; Cat# 950–223) at 100 °C for 24 min to disrupt the primary antibody–enzyme complex and minimise antibody carryover, thereby preventing binding of subsequent chromogens to residual enzyme activity.

The AT8/GFAP and pTau217/GFAP two-plex stainings were performed by first incubating anti-AT8 or anti-pTau217 for 60 min, followed by detection with OmniMap anti-mouse HRP or with OmniMap anti-rabbit HRP for 16 min and visualisation using the Discovery Purple Kit. Anti-GFAP was then applied for 60 min, detected using OmniMap anti-rabbit HRP for 16 min and developed using Discovery Green HRP. The AT8/NeuN two-plex staining was initiated by incubating anti-AT8 for 60 min, followed by detection using Discovery UltraMap anti-mouse HRP and visualisation using the Discovery Purple Kit. Subsequently, anti-NeuN was applied for 60 min, detected using OmniMap anti-mouse HRP, and developed using Discovery Teal HRP. The pS422/NeuN and pS422/VGAT two-plex stainings were performed by incubating anti-pS422 for 60 min, followed by detection using Discovery UltraMap anti-rabbit HRP and visualisation using the Discovery Red Kit. Subsequently, anti-NeuN or anti-VGAT was applied for 60 min, detected using OmniMap anti-mouse HRP or OmniMap anti-rabbit HRP, and developed using Discovery Teal HRP.

The SST/AT8 and PVALB/AT8 two-plex stainings were initiated by incubating anti-SST or anti-PVALB for 60 min, followed by detection using OmniMap anti-rabbit HRP for 16 min and visualisation using Discovery Teal HRP. Subsequently, anti-AT8 was applied for 60 min, detected using Discovery UltraMap anti-mouse AP and developed using the Discovery Yellow Kit.

For the SST/pS396 and PVALB/pS396 two-plex panels, anti-SST or anti-PVALB were first incubated for 60 min, detected using OmniMap anti-rabbit HRP for 16 min and visualised using the Discovery CM DAB kit. Anti-pS396 was subsequently applied for 60 min, detected using OmniMap anti-rabbit HRP for 16 min and developed using Discovery Teal HRP.

The pTau217/AT8, pTau217/GT38, pTau217/pS396, pS396/AT8, pS396/GT38 and GT38/AT8 two-plex stainings were initiated by incubating anti-pTau217, anti-pS396 or anti-GT38 for 60 min, followed by detection using OmniMap anti-rabbit HRP or OmniMap anti-mouse HRP for 16 min and visualisation using the Discovery Purple Kit. Anti-AT8, anti-GT38 or anti-pS396 were then applied for 60 min, detected using OmniMap anti-mouse or anti-rabbit HRP for 16 min, as appropriate, and developed using Discovery Teal HRP.

For the GT38/4G8 and 4G8/AT8 two-plex panels, anti-GT38 or anti-4G8 was first incubated for 60 min, detected using OmniMap anti-mouse HRP for 16 min and developed using Discovery Teal HRP. Anti-4G8 or anti-AT8 was subsequently applied for 60 min, detected using OmniMap anti-mouse HRP for 16 min and visualised using the Discovery Purple Kit. For the pS422/4G8 and pTau217/4G8 two-plex panels, anti- pS422 or anti-pTau217 was first incubated for 60 min, detected using OmniMap anti-rabbit HRP for 16 min and visualised using the Discovery Red Kit. Anti-4G8 was subsequently applied for 60 min, detected using OmniMap anti-mouse HRP for 16 min and developed using Discovery Teal HRP. For the pS396/4G8 two-plex panel, anti- pS396 was first incubated for 60 min, detected using OmniMap anti-rabbit HRP for 16 min and developed using Discovery Teal HRP. Anti-4G8 was subsequently applied for 60 min, detected using OmniMap anti-mouse HRP for 16 min and visualised using the Discovery Purple Kit.

The pS422/pSyn81A two-plex staining was initiated by incubating anti-pS422 for 60 min, followed by detection with Discovery UltraMap anti-rabbit AP and visualisation using the Discovery Red chromogen kit. Anti-pSyn81A was then applied for 60 min, detected using OmniMap anti-rabbit HRP for 16 min and developed using Discovery Teal HRP.

For the pSyn81A/pS396, pSyn81A/pTau217, pSyn81A/AT8 and pSyn81A/GT38 two-plex stainings, anti-pSyn81A was first incubated for 60 min, detected using OmniMap anti-rabbit HRP for 16 min and visualised using the Discovery CM DAB kit. Anti-pS396, anti-pTau217, anti-AT8 or anti-GT38 were subsequently applied for 60 min, detected using OmniMap anti-rabbit or anti-mouse HRP for 16 min, as appropriate, and developed using Discovery Teal HRP.

The three-plex stainings pS422/pS396/GT38, pS422/pS396/AT8 and pS422/pS396/pTau217 were performed by first incubating anti-pS422 for 60 min, followed by detection with Discovery UltraMap anti-rabbit AP and visualisation using the Discovery Red chromogen kit. In a second step, anti-pS396 was applied for 60 min, detected using OmniMap anti-rabbit HRP for 16 min and developed using Discovery Teal HRP. Finally, GT38, AT8 or pTau217 were incubated for 60 min, detected using OmniMap anti-mouse HRP for 16 min and visualised using the Discovery Green HRP kit (Cat# 760–271).

For the pTau217/pS396/AT8 three-plex panel, anti-pTau217 was first incubated for 60 min, detected using Discovery UltraMap anti-rabbit AP and developed with the Discovery Red chromogen kit. Anti-pS396 was subsequently applied for 60 min, detected using OmniMap anti-rabbit HRP for 16 min and visualised using Discovery Teal HRP. Finally, anti-AT8 was incubated for 60 min, detected using OmniMap anti-mouse HRP for 16 min and developed using Discovery Green HRP.

The GT38/pS396/AT8 three-plex staining was initiated by incubating anti-GT38 for 60 min, followed by detection with OmniMap anti-mouse HRP for 16 min and visualisation using the Discovery CM DAB kit. Subsequently, anti-pS396 was applied for 60 min, detected using OmniMap anti-rabbit HRP for 16 min and developed using the Discovery Purple Kit. Finally, anti-AT8 was incubated for 60 min, detected using OmniMap anti-mouse HRP for 16 min and visualised using Discovery Teal HRP.

The four-plex pSyn81A/pS396/4G8/AT8 staining was initiated by incubating sections with anti-pSyn81A for 60 min, followed by detection with OmniMap anti-rabbit HRP for 16 min and visualisation using the Discovery CM DAB kit. Subsequently, anti-pS396 was applied for 60 min, detected with OmniMap anti-rabbit HRP for 16 min and developed using the Discovery Purple Kit (Cat# 760–229). Anti-4G8 was then incubated for 60 min, detected with Discovery UltraMap anti-mouse alkaline phosphatase (AP) and visualised using the Discovery Yellow Kit (Cat# 760–239). Finally, anti-AT8 was applied for 60 min, detected with OmniMap anti-mouse HRP for 16 min and developed using the Discovery Teal HRP kit (Cat# 760–247).

Upon completion of the multiplex staining procedure, slides were counterstained with haematoxylin, dehydrated through graded ethanol and xylene, and coverslipped following the same protocol used for single-target IHC using the automated slide coverslipper Tissue-Tek Prisma® Plus and Tissue-Tek Film® (Sakura Finetek).

### 2.6 Multiplex immunohistofluorescence (mIHF)

Multiplex IF was performed on the same automated platform. Following deparaffinisation, tissue sections underwent heat-induced epitope retrieval (HIER) in CC1 at 95 °C for 64 min. Endogenous peroxidase activity and nonspecific background staining were suppressed by incubation with Discovery Inhibitor for 8 min at 37 °C. Primary antibodies targeting AT8, pS396, pS422, pSyn81A, GT38 and pTau217 were diluted in EnVision™ FLEX antibody diluent (**Supplementary Table 2**) and manually applied for 60 min. Detection was performed using species-appropriate HRP-conjugated OmniMap secondary reagents (anti-Mouse or anti-Rabbit) for 12 min, followed by tyramide signal amplification using tyramide-conjugated fluorophores applied for 8 min. Fluorophores included Rhodamine 6G (R6G, Cat# 760-233, λ_ex ≈ 542 nm / λ_em ≈ 568 nm), Cyanine 5 (Cy5, Cat# 760-238, λ_ex ≈ 650 nm / λ_em ≈ 670 nm) and fluorescein amidite (FAM, Cat# 760-243, λ_ex ≈ 490 nm / λ_em ≈ 520 nm). Between sequential labelling steps, heat-mediated deactivation (HD; 100 °C, 24 min) using CC2 was performed to minimise cross-reactivity and ensure signal specificity.

Four duplex stainings were performed using AT8 and either pS396, pS422, pSyn81A or GT38. In all panels, AT8 was developed using Cy5, while pS396, pS422, pSyn81A and GT38 were visualised using FAM. For the pS422/pS396 duplex panel, anti-pS422 was first applied for 60 min, detected with species-appropriate OmniMap HRP secondary reagents and developed using Rhodamine 6G. Anti-pS396 was subsequently incubated for 60 min and visualised using Cy5. Finally, a duplex pTau217/AT8 staining was performed, with pTau217 developed using Rhodamine 6G and AT8 visualised using Cy5.

The GT38/AT8/pS396, pTau217/AT8/pS396, pSyn81A/AT8/4G8 and pTau217/AT8/GFAP three-plex panels were initiated by incubating anti-GT38, anti- pTau217 or anti-pSyn81A for 60 min, followed by detection with species-appropriate OmniMap HRP secondary reagents and development using Rhodamine 6G. Subsequently, anti-AT8 was applied for 60 min and visualised with Cy5. Finally, pS396, anti-4G8 or anti-GFAP was incubated for 60 min and developed using FAM. A duplex GT38/pS396 staining was also performed, with GT38 developed using Rhodamine 6G and pS396 visualised using FAM.

Upon completion of fluorescent labelling, slides were rinsed in dH₂O, air-dried at room temperature for 10 min while protected from light, and manually coverslipped using ProLong™ Gold Antifade Mountant (Cat# P36934; Thermo Fisher Scientific) to preserve fluorescence signals.

### 2.7 Image acquisition and processing

Chromogenically stained FFPE tissue sections were acquired by brightfield microscopy using a Leica DM2000 LED microscope equipped with a range of objectives (5×, 10×, 20×, 40×, and 63×) and a LMS Leica Flexacam i5 camera. For larger-scale visualization and marker quantification on the HALO® image analysis platform (v4.0.5107.445, Indica Labs, Albuquerque, NM, USA), entire tissue sections were captured through high-throughput whole-slide scanning using the IntelliSite Ultra Fast Scanner (Philips). For specific analyses requiring detailed quantification, images were also exported directly from the HALO® image analysis platform. Fluorescently labelled sections were imaged in three dimensions by confocal microscopy. Z-stack images were acquired using Zeiss LSM800 confocal microscopes equipped with 20× air (Plan-Apochromat 20×/0.8) and 40× oil immersion (EC Plan-Neofluar 40×/1.3 Oil DIC) objectives. Three-dimensional image datasets were processed, visualised, and analysed in Imaris software (version 11.0.1). Channel-specific intensity normalisation and background subtraction were applied to reduce noise and improve signal fidelity, allowing accurate visualisation and assessment of signal distribution and spatial relationships within the tissue.

### 2.8 Dataset

Publicly available hippocampal microarray data were obtained from the Gene Expression Omnibus (GEO accession GSE48350; Affymetrix HG-U133 Plus 2.0 platform). The dataset comprised 43 non-demented control samples and 19 AD samples, classified according to the diagnostic labels provided by the original submitters. Raw expression values were log₂-transformed prior to all analyses. Two gene panels were defined a priori based on established cell-type marker specificity: general neuronal marker (*RBFOX3*), a GABAergic/interneuron panel (*SST*, *GAD1*, *GAD2*, *SLC32A1*) and an excitatory neuronal panel (*SLC17A7*, *CAMK2A*, *GRIN1*, *GRIA2*). For each gene individually, group differences between AD and control were assessed using two-sided Welch’s t-tests on log₂ expression values, with mean fold-changes back-transformed to linear scale for reporting.

## 3 Results

### 3.1 Cohort characterisation and disease-specific co-pathology of CTL, AD, PDD and DLB hippocampal samples

Demographic and neuropathological characteristics of cohort 1 (CTL, AD, PDD) are summarised in **Fig. 1C**. Groups were comparable in gender distribution and *post-mortem* delay. Mean age at death was slightly lower in both disease conditions, AD and PDD, compared to CTL. Braak NFT stage ranged from II–IV in CTL, IV–VI in AD, and 0-II in PDD (**Fig. 1C**). Several CTL cases additionally showed astrocytic tau pathology consistent with age-related tau astrogliopathy (ARTAG) (**Supplementary Fig. S1; Table 1**).

To characterise the co-pathology landscape of cohort 1, we quantified AT8 immunoreactivity as a reference marker of tau pathology alongside 4G8 for amyloid-β and pSyn81A for α-synuclein (**Fig. 1D**). At the whole-hippocampus level, AT8 immunoreactivity was significantly higher in AD than in both CTL and PDD (p<0.01 for both comparisons. AT8 labels phosphorylated tau across the spectrum of tau pathology, including pretangles, mature tangles, ghost tangles, neuropil threads, neuritic plaques and tangle-associated neuritic clusters (TANCs) (**Fig. 1E**). 4G8 immunoreactivity was significantly higher in AD than in both CTL and PDD (p<0.0001 for both comparisons), with AD cases showing diffuse plaques, neuritic plaques, and cerebral amyloid angiopathy (CAA) across all subfields (**Fig. 1F, Supplementary Fig. S2A**). CTL cases showed low but detectable 4G8 immunoreactivity (mean coverage 0.02%) (**Fig. 1D**). PDD showed similarly low 4G8 levels (**Fig. 1D**). pSyn81A immunoreactivity was significantly higher in PDD than in both CTL and AD (p<0.0001 for both comparisons) (**Fig. 1D**), with Lewy body, Lewy neurite, and tangle-like α- synuclein morphology distributed across subfields and highest burden in CA2 (**Fig. 1G, Supplementary Fig. S2B**). CTL cases showed low but detectable pSyn81A immunoreactivity (mean coverage 0.02%) (**Fig. 1D**). AD cases showed very low pSyn81A signal (mean coverage 0.001%) (**Fig. 1D**, **Supplementary Fig. S2B**).

Together, these observations indicate that cohort 1 is stratified by a co-pathology profile distinct to each diagnostic group. To extend this comparison, we examined a second, independent cohort of CTL, AD, and DLB cases from a different brain bank (cohort 2; **Supplementary Fig. S3A-C**), which differed from cohort 1 in gender distribution and PMD. Quantification of AT8, 4G8, and pSyn81A in cohort 2 showed co-pathology profile in DLB more similar to AD across hippocampal subfields (**Supplementary Fig. S3D–F**). A direct comparison between DLB and PDD cases across the two cohorts was not attempted given the differences in sample processing between brain bank sources.

### 3.2 Divergent phospho-tau epitope profiles distinguish AD and PDD hippocampal pathology

To determine the characteristics of tau pathology across disease conditions, markers were selected to cover early phosphorylation (pS422), broad hyperphosphorylation (AT8), mature phosphorylation (pS396), biomarker-relevant late epitope (pTau217) and conformational change (GT38), providing a multi-dimensional readout of tangle composition across the maturation sequence [10,12,22,23]. All markers were tested across a set of CTL, AD, PDD, DLB, PSP and 3R-tau FTLD samples validating the immunoreactivity of tangles across diseases (**Supplementary Fig. S4**). The regional distribution of the DAB-stained area of five tau markers was analysed across six hippocampal subfields (DG, CA4, CA3, CA2, CA1 and subiculum) and the parahippocampal cortex (PHC) in non-demented CTL (n=9), AD (n=13) and PDD (n=14) cases of cohort 1.

AT8, pTau217, and GT38 immunoreactivity was consistently higher in AD than in CTL and PDD across most of the examined subfields and in PHC (all comparisons AD vs. PDD, p<0.05 to p<0.0001; **Fig. 2A-B, C-D, E-F**). For each of these three markers the PDD DAB-stained area did not exceed the levels of CTL and was in several regions lower than CTL. In contrast, pS396 immunoreactivity followed a distinct pattern. pS396 burden was significantly enriched in AD compared to CTL (all subfields, p<0.05 to p<0.0001) but PDD showed higher but more variable numerical value in most subfields to the exception of the subiculum and PHC (**Fig. 2G-H**). pS422 displayed a distinct pattern, with significantly higher DAB-stained area in PDD than AD in total hippocampus, and across CA2, CA1, subiculum, and PHC (p<0.02 to p<0.001). In addition, pS422 burden was increased in PDD compared with CTL in CA3 and CA2 (p<0.04, **Fig. 2I-J**). Absolute pS422 burden was low (<1% stained area) compared to the other markers across all groups and regions.

**Figure 2.**
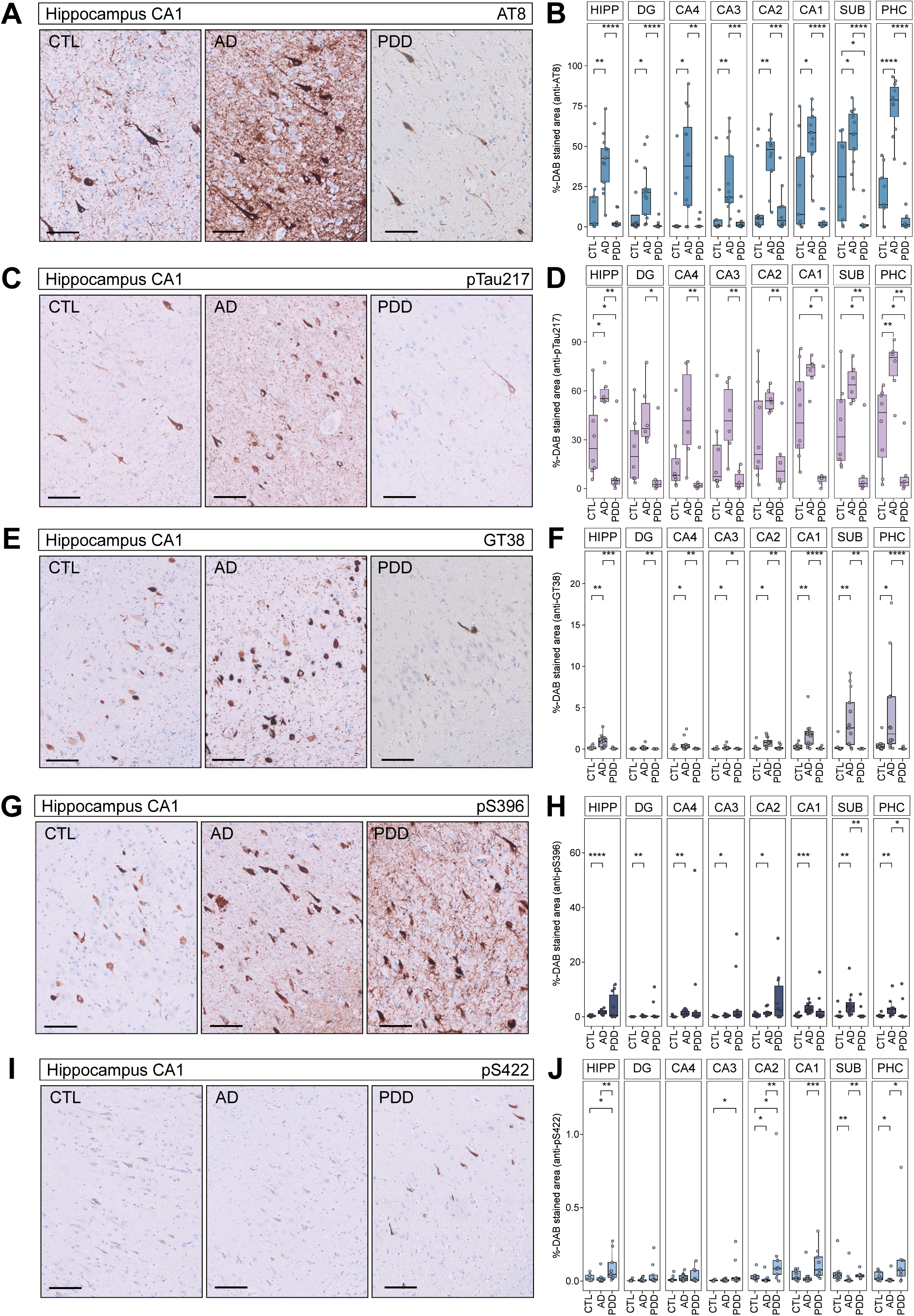
Divergent phospho-tau and conformational immunophenotypes distinguish AD and PDD hippocampal and parahippocampal pathology. Representative chromogenic immunohistochemistry images and staining quantification (%-DAB stained area) in dentate gyrus (DG), CA4, CA3, CA2, CA1, subiculum (SUB), parahippocampal cortex (PHC), and combined hippocampal subfields (HIPP: sum of DG, CA4, CA3, CA2, and CA1) of **A, B** anti-AT8 (pS202/pT205) (**A** case#2, #31, #47); **C, D** anti-pTau217 (**C** case#5, #32, #49); **E, F** conformation-selective tau antibody GT38 (**E** case#6, #31, #49); **G, H** anti- pS396 (**G** case#7, #32, #53) and **I, J** anti-pS422 (**I** case#4, #20, #49) in non-demented controls (CTL), Alzheimer’s disease (AD), and Parkinson’s disease dementia (PDD). Boxplots show median and interquartile range and each point represents one case. Statistical comparisons were performed by Wilcoxon rank-sum test, with statistical significance set at p<0.05. Levels of significance are indicated as follows: p<0.05 (*), p<0.005 (**), and p<0.0005 (***), p<0.00005 (****). Scale bars: **A**, **C**, **E**, **G**, **I** 100 µm.

To determine whether the tau epitope differences observed between AD and PDD in cohort 1 extended to AD and DLB, we quantified the five tau markers across hippocampal subfields and PHC in cohort 2 (non-demented CTL (n=9), AD (n=11) and DLB (n=14); **Supplementary Fig. S3F, S5 A-D**). AT8, pTau217, GT38, and pS396 DAB-stained areas were significantly elevated in both AD and DLB relative to CTL across most subfields, with no significant difference between AD and DLB for any of these markers (**Supplementary Fig. S3F, S5C**). The pS422 burden showed little difference between groups, except for PHC, where DLB DAB-stained area was significantly higher than in AD (**Supplementary Fig. S5D**).

Qualitative assessment across hippocampal subfields and PHC showed that the five tau markers labelled overlapping but distinct pathological morphologies, including pretangles, mature and ghost tangles, neuropil threads, neuritic plaques, and tangle-associated neuritic clusters (TANCs), with pS396 and pTau217 showing a more restricted morphological staining in PDD than in CTL or AD (**Supplementary Fig. S6A**). Total-hippocampal tau marker burden did not show association with Braak staging within any diagnostic group, although several markers showed non-significant trends, including a negative trend for pS422 in AD (R=−0.5, p=0.099) and a positive trend for pS396 in PDD (R=0.57, p=0.086) (**Supplementary Fig. S6B-F**).

### 3.3 Tau epitope covariation differs by diagnostic group

We performed pairwise Spearman correlation of the regional DAB-stained area for AT8, pS396, pS422, pTau217, and GT38, to determine whether the regional and subregional burden of individual tau epitopes covaried within diagnostic group (non-demented CTL (n=9), AD (n=12) and PDD (n=10); cohort 1).

In CTL hippocampus, all five markers showed positive correlations with one another (**Fig. 3A**). This pattern was altered in AD where AT8/GT38 and pS396/GT38 remained positively correlated with each other at the whole-hippocampus level (**Fig. 3B**). This AT8/GT38 and pS396/GT38 correlation are partially presented at the subfield level. In DG, CA4 and CA3, we observe significant pairwise positive correlation of AT8/GT38 and pS396/GT38 (AT8/GT38: DG r=0.77, p=0.003; CA4 r=0.76, p=0.006; CA3 r=0.83, p=0.002; ps396/GT38: DG r=0.87, p=0.001; CA4 r=0.85, p=0.002; CA3 r=0.82, p=0.007), while DG and CA3 present additional significant positive correlation between AT8 and ps396 (AT8/pS396: DG r=0.65, p=0.034; CA3 r=0.72, p=0.017) (**Supplementary Fig. S7A-C**). AD CA2, CA1 and PHC only present a significant positive correlation between pS396 and GT38 (pS396/GT38: CA2 r=0.64, p=0.04; CA1 r=0.68, p=0.025; PHC r=0.89, p=0.000) (**Supplementary Fig. S7D-F**). In PHC

**Figure 3.**
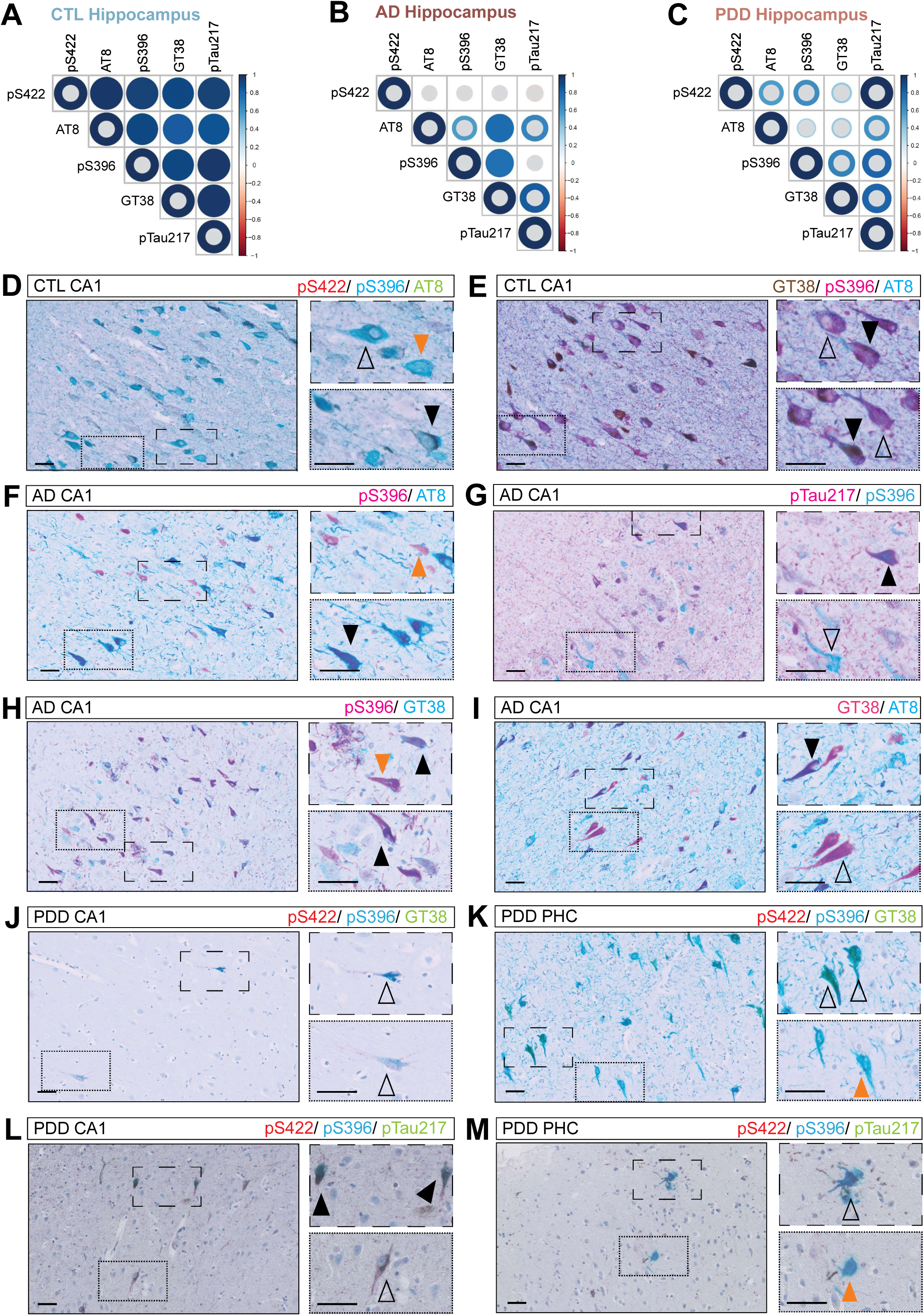
Tau epitope covariation and diagnosis-specific multiplex co-positivity. Pairwise Spearman correlation matrix of regional tau marker burden (pS422, AT8, pS396, GT38, pTau217) in **A** CTL hippocampus (n=9), **B** AD (n=12) and **C** PDD (n=10). Each cell reports the Spearman correlation coefficient (ρ) between %DAB-stained area for the two indicated tau markers, calculated across all cases within the diagnostic group. Colour scale indicates the direction and magnitude of ρ (blue = positive, red = negative, range 1 to −1). Correlations were considered significant at p<0.05 (two-sided). Non-significant correlations are marked with a grey dot. *P* values were obtained from two-sided Spearman rank correlation tests performed separately for each pair of markers. **D** Representative triplex cIHC of pS422(red)/pS396(teal)/AT8 (green) (case#9) and **E** GT38(DAB)/pS396(purple)/AT8(teal) in the CTL CA1, illustrating co-occurrence of multiple phospho-tau epitopes within individual tangles (case#9). **F–I** Representative duplex cIHC for pS396 (purple or teal), pTau217 (purple), AT8 (teal) and GT38 (teal) combinations in the AD CA1, illustrating distinct tangle subpopulations by differential marker co-positivity (case#45, #32, #46, #44). Representative pS422(red)/pS396(teal)/GT38(green) cIHC of PDD **J** CA1/CA2 and **K** PHC (case#47). Representative pS422(red)/pS396(teal)/pTau217(green) cIHC of PDD **L** CA1/CA2 and **M** PHC (case#60). Back arrowheads indicate single-staining, while transparent arrowheads indicate double-/triple-positive. Orange arrowheads indicate pS396 single-positive tangles across several staining combination and disease-conditions. Scale bars: **D-M** 50 µm.

AT8 showed an additional significant, but negative correlation with pS422 (r=−0.64, p=0.04) (**Supplementary Fig. S7F**). In PDD, none of the tau markers significantly correlated at the whole-hippocampus level (**Fig. 3C**). This observation was not uniform across subfields. In DG, CA4 and CA3, no pairwise correlation reached statistical significance (**Supplementary Fig. S7G-I**). In CA2, pS422 formed a significantly correlated set together with pS396 and GT38 (pS422/pS396 r=0.88, p=0.003; pS422/GT38 r=0.79, p=0.028; pS396/GT38 r=0.88, p=0.003) (**Supplementary Fig. S7J**). In CA1, AT8, pS396, and GT38 showed significant correlation (AT8/pS396 r=0.68, p=0.035; AT8/GT38 r=0.70, p=0.043; pS396/GT38 r=0.68, p=0.05) (**Supplementary Fig. S7K**). In PDD PHC, a further distinct pattern emerged: AT8 and GT38 showed a significant positive correlation (r=0.88, p=0.007) (**Supplementary Fig. S7L**).

### 3.4 Multiplex co-staining reveals diagnosis-specific tangle marker co-positivity

Non-demented CTL cases showed a consistent pattern of tau immunoreactivity across all markers in the hippocampus. We therefore used multiplex cIHC to define single- tangle epitope signatures. In CTL hippocampus, triplex cIHC for pS422, pS396, and AT8/GT38 showed multiple marker combinations within individual tangles. Single-positive pS396 tangles were observed alongside double-positive pS422/pS396 and triple-positive tangles, indicating that multiple phospho-tau epitopes can co-occur within the same tangle even in non-demented condition (**Fig. 3D; Supplementary Fig. S8A**). Even though GT38 staining was relatively sparse in CTL, triple-positive tangles and double-positive pS396/AT8 were common in GT38/pS396/AT8 co-stained sections, whereas AT8 and GT38 single-positive tangles were essentially absent (**Fig. 3E**). pTau217 immunoreactivity was observed predominantly within double-positive pS422 or pS396, or triple-positive tangles with AT8 (**Supplementary Fig. S8B-C**). In a subset of CTL cases, we found astrocytic tau pathology that was confirmed by co-staining with the astrocyte marker GFAP alongside AT8 and pTau217 by chromogenic duplex immunohistochemistry (**Supplementary Fig. S8 D, E**) and by triplex immunofluorescence (**Supplementary Fig. S8F**), demonstrating colocalisation of both tau epitopes within GFAP-positive astrocytic processes.

In AD, different duplex cIHC for pS396, AT8, GT38 and pTau217 identified distinct tangle subpopulations distinguished by differential marker co-positivity (**Fig. 3F-I; Supplementary Fig. S8G, H**). Across several marker combination, we observe single-positive tangles beside double-positive tangles within the same hippocampal subfield. Notably, AT8- and pS396 single-positive tangles were observed across every duplex combination examined, regardless with which second marker it was paired. In contrast, pTau217 immunoreactivity was consistently detected only in combination with at least one other marker (**Fig. 3F-I; Supplementary Fig. S8G, H**).

In PDD, sections stained with pS422/pS396/GT38 showed that pS422 immunoreactivity in CA1 and CA2 was predominantly neuritic. pS422/pS396 double-positive tangles were the most common. In the PHC, pS396 single-positive and pS396/GT38 double-positive tangles were observed in approximately equal proportion (**Fig. 3J, K**). Triplex staining of pS422/pS396/pTau217 showed triple-positive tangles in CA1 and PHC (**Fig. 3L, M**).

In DLB, GT38/pS396/AT8 triplex co-staining revealed heterogeneous tangle immunophenotypes across all hippocampal subfields examined, with single-positive tangles observed alongside double- and triple-positive tangles within the same subfield, comparable to the co-positivity pattern described above in AD (**Supplementary Fig. S9A**).

### 3.5 Single-tangle classification reveals regionally distinct phospho-tau/conformational composition in AD and PDD

To address how tau epitope heterogeneity is organised across the hippocampal subfield neuronal populations in AD and PDD, we classified individual tangles after triplex cIHC AT8/pS396/GT38 into one of seven possible combinations across DG, CA4, CA3, CA2, CA1, and subiculum in AD and PDD cases (n=3; AD: 4.737 tangles analysed in AD and 347 tangles analysed in PDD).

In AD, two immunophenotypes showed a graded change in their relative abundance across the hippocampal subfield sequence from DG to subiculum. Tangles positive for pS396 and GT38 but negative for AT8 (pS396+/AT8−/GT38+, orange) comprised 22.2% of classified tangles in DG, declining to 12.0% in CA1 and 7.2% in subiculum (**Fig. 4A, B**). Conversely, tangles positive for AT8 alone (pS396−/AT8+/GT38−, teal) comprised 17.6% of tangles in DG, increasing to 34.6% in CA1 and 50.2% in subiculum (**Fig. 4A, B**). Triple-positive tangles (pS396+/AT8+/GT38+, black) remained comparatively stable across DG through CA1 (11.3–15.0%) and were lower in subiculum (6.6%) (**Fig. 4A, B**). This gradient was not observed across all hippocampal subfields. CA3 showed the highest proportion of pS396+/AT8−/GT38+ tangles (37.0%) and the lowest proportion of AT8-only tangles (10.9%) of any region examined (**Fig. 4A-B**). To be noted, tangle counts in DG and CA3 were low and unevenly distributed across the three cases.

**Figure 4.**
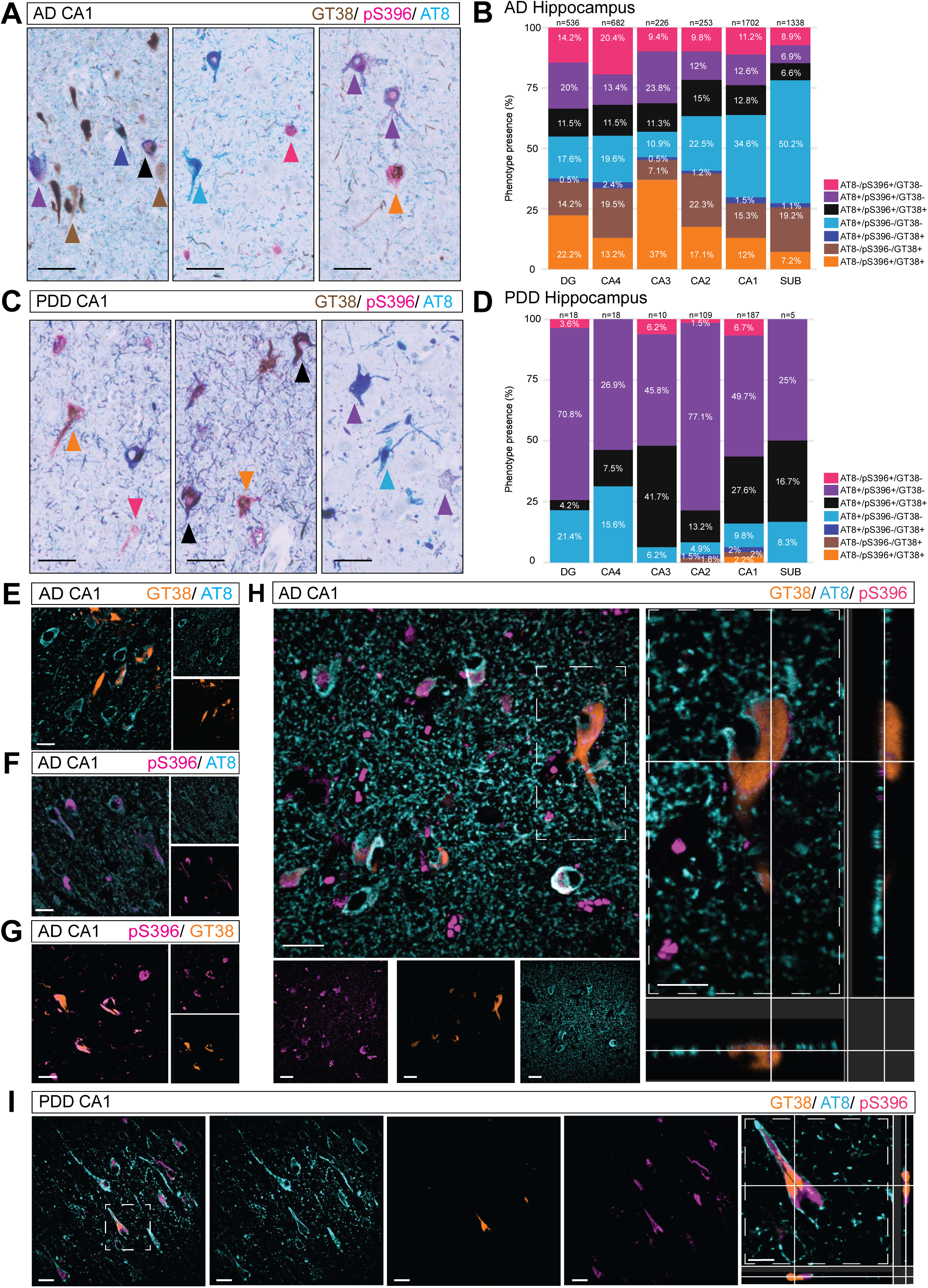
Single-tangle classification reveals regionally graded phospho- tau/conformational composition in AD. **A** Representative chromogenic multiplex images illustrating the seven possible GT38(DAB)/pS396(purple)/AT8(teal) single-tangle immunophenotypes in AD hippocampus (case#31). Arrowheads indicate marker-positive tangles according to the tangle immunophenotype legend. **B** Mean percentage of single tangles classified into each immunophenotype across DG, CA4, CA3, CA2, CA1, and subiculum (n=3 cases). Percentages are reported as the mean across the three cases, with each case weighted equally regardless of tangle number. **C** Representative chromogenic multiplex images illustrating the seven possible GT38(DAB)/pS396(purple)/AT8(teal) single-tangle immunophenotypes in PDD hippocampus (case #47). Arrowheads indicate marker-positive tangles according to the tangle immunophenotype legend. **D** Mean percentage of single tangles classified into each immunophenotype across DG, CA4, CA3, CA2, CA1, and subiculum, PDD (n=3 cases). Percentages are reported as the mean across the three cases, with each case weighted equally regardless of tangle number. **E–G** Representative duplex immunofluorescence for pS396 (purple), AT8 (teal), and GT38 (orange) combinations in AD CA1, illustrating distinct tangle subpopulations by differential marker co-positivity (cases #27, #20, #42). **H-I** High- resolution confocal z-stack imaging of GT38(orange)/AT8(teal)/pS396(purple) triplex- stained **H** AD CA1 and **I** PDD CA1 showing subcellular NFT organisation: AT8 at the outer rim, pS396 in an inner compartment, GT38 innermost (case#20, #61). Scale bars: **A**, **C** 50 µm; **E–G** 10 µm; **H, I** 30 µm (overview), 15 µm (inset).

Individual tangle epitope classification was additionally performed on PDD, though overall tangle counts were substantially lower than in AD across all regions (**Fig. 4C, D**). The AT8+/pS396+/GT38− immunophenotype was the most common profile in every region examined, ranging from 26.9% in CA3 and 25.0% in subiculum to 77.1% in CA2. Triple-positive (AT8+/pS396+/GT38+, black) tangles were the second most common immunophenotype in most regions, comprising 4.2% in DG, 7.5% in CA4, 41.7% in CA3, 13.2% in CA2, 27.6% in CA1, and 16.7% in subiculum. GT38-positive tangles other than the triple-positive tangles were rare, together comprising less than 10% of classified tangles in any region (**Fig. 4C, D**).

### 3.6 Intra-tangle spatial architecture of phospho-tau and conformational epitopes

In AD, duplex IF for pS396, AT8 and GT38 identified distinct mature tangle subpopulations distinguished by differential marker co-positivity, with AT8 immunoreactivity consistently localised to the tangle periphery relative to pS396 or GT38, which occupied more central regions of the same tangles (**Fig. 4E-G**). To confirm and further resolve this spatial relationship at higher resolution, we examined morphologically mature NFTs in CA1 by high-resolution confocal z-stack imaging of AT8/pS396/GT38 and AT8/pS396/pTau217 triplex-stained sections in AD (**Fig. 4H; Supplementary Fig. S9B**). These multiplex cIHC showed the same rim-and-core spatial organisation observed by duplex staining: AT8 immunoreactivity was localized to the outer rim of the tangle, delineating its perimeter, while pS396 and pTau217 occupied an inner tangle compartment. Triplex imaging additionally resolved GT38 as confined to the innermost compartment, a distinction not resolvable by duplex staining alone (**Fig. 4H**).

In PDD, AT8 and pS396 showed a comparable rim-and-core spatial relationship to that observed in AD, in both hippocampus and PHC (**Fig. 4I**). In the few tangles where GT38 was present, it occupied the same innermost compartment as in AD (**Fig. 4I**). In a DLB case, the confocal appearance of AT8, pS396, and GT38 closely resembled the AD pattern described above (**Supplementary Fig. S9C**), further suggesting that this subcellular organisation is stereotypical across disorders.

### 3.7 Neuronal and synaptic marker density shows regionally distinct, disease-associated patterns in AD and PDD

To characterise the hippocampal neuronal and synaptic composition in relation to tau, amyloid, and α-synuclein pathology, we quantified five markers reflecting distinct neuronal compartments across six hippocampal subfields, including subiculum, and PHC in CTL, AD, and PDD: NeuN (overall neuronal density), vesicular GABA transporter (VGAT, inhibitory presynaptic terminals), vesicular glutamate transporter 1 (VGLUT1, excitatory presynaptic terminals), and the interneuron populations marked by somatostatin (SST) and parvalbumin (PVALB).

NeuN immunoreactivity showed a declining trend in both AD and PDD relative to CTL that did not reach statistical significance at the whole-hippocampus level (CTL vs. AD, p=0.67; CTL vs. PDD, p=0.76; AD vs. PDD, p=0.63). This pattern was not uniform across subfields: in DG, CA4, CA3, and PHC, NeuN immunoreactivity in PDD remained close to CTL levels while AD showed a numerically greater reduction. In CA2, CA1, and subiculum, AD and PDD showed comparable reductions relative to CTL, reaching significance only in CA1 between AD and PDD (**Fig. 5A**). VGAT immunoreactivity showed no significant difference between groups at the whole-hippocampus level (CTL vs. AD, p=0.43; CTL vs. PDD, p=0.22; AD vs. PDD, p=0.51) or in any individual subfield (**Fig. 5B**). VGLUT1 immunoreactivity was lower in both AD and PDD relative to CTL across multiple subfields, reaching significance only in CA2 (AD, p=0.04; PDD, p=0.02) and at the whole-hippocampus level (AD, p<0.05; PDD, p<0.05). No significant difference was detected between AD and PDD in any subfield (**Fig. 5C**). SST and PVALB immunoreactivity were comparable across groups in most subfields (**Supplementary Fig. S10**).

**Figure 5.**
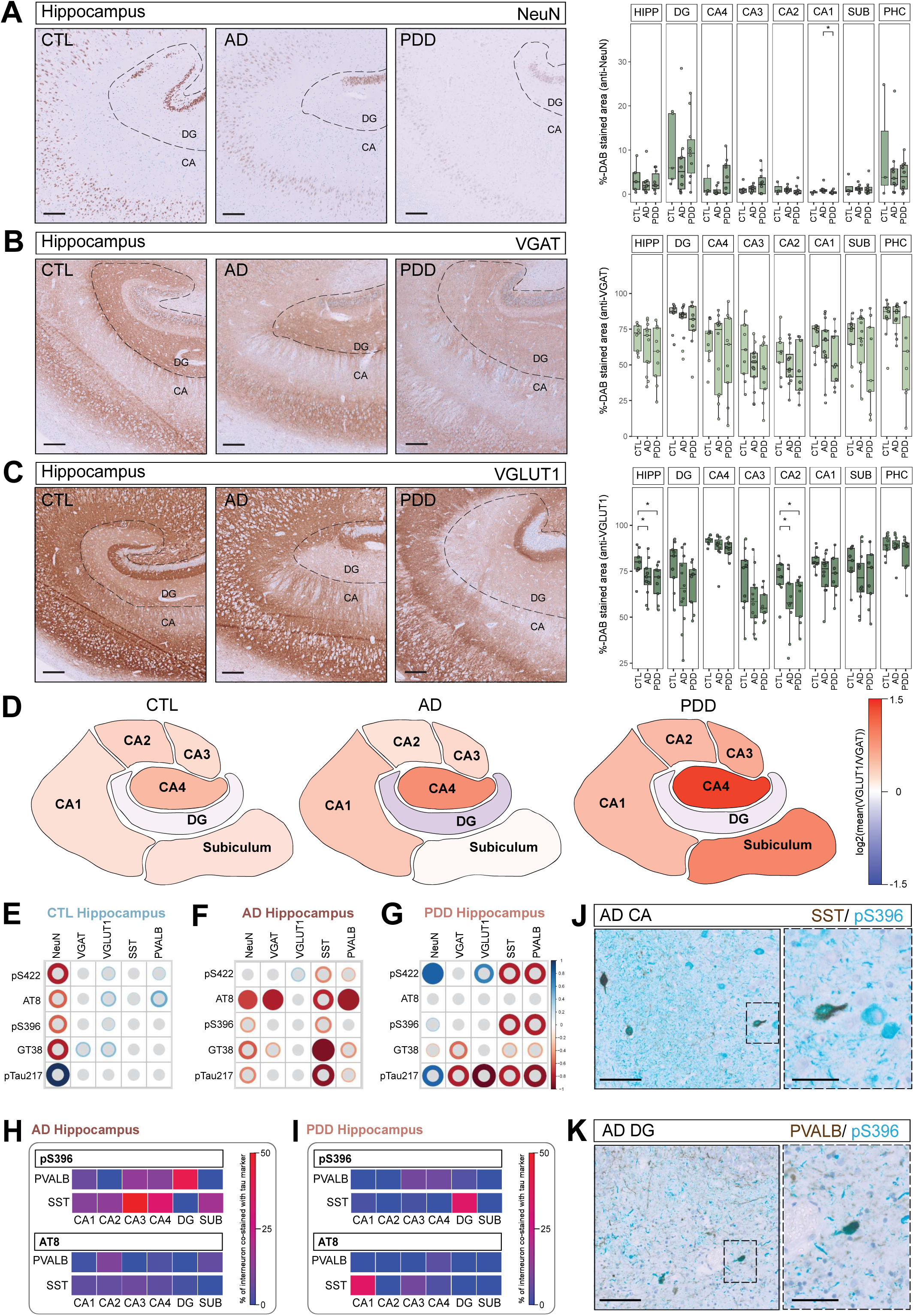
Neuronal and synaptic marker density and its relationship to tau burden in AD and PDD. Representative chromogenic immunohistochemistry images and staining quantification (%-DAB stained area) in dentate gyrus (DG), CA4, CA3, CA2, CA1, subiculum (SUB), parahippocampal cortex (PHC), and combined hippocampal subfields (HIPP: sum of DG, CA4, CA3, CA2, and CA1) of **A** NeuN (case#9, #32, #47), **B** VGAT (case#9, #32, #47) and **C** VGLUT1 (case#9, #32, #47). Boxplots show median and interquartile range and each point represents one case. Statistical comparisons were performed by Wilcoxon rank-sum test, with statistical significance set at p<0.05. Levels of significance are indicated as follows: p<0.05 (*), p<0.005 (**), and p<0.0005 (***), p<0.00005 (****). **D** Regional log2(VGLUT1/VGAT) immunoreactivity ratio by subfield and diagnostic group. **E** Pairwise Spearman correlation matrix, five tau markers with five neuronal/synaptic markers (NeuN, VGAT, VGLUT1, SST, PVALB) in CTL, **F** AD and **G** PDD hippocampus. Each cell reports the Spearman correlation coefficient (ρ) between %DAB-stained area for the two indicated tau markers, calculated across all cases within the diagnostic group. Colour scale indicates the direction and magnitude of ρ (blue= positive, red= negative, range 1 to − 1). Correlations were considered significant at p<0.05 (two-sided). Non-significant correlations are marked with a grey dot. *P* values were obtained from two-sided Spearman rank correlation tests performed separately for each pair of markers. **H, I** Heatmap of percentage SST/PVALB interneuron co-positivity with AT8 or pS396 across subfields in AD (**H**, n=3) and in PDD (**I**, n=3). **J** Representative SST(DAB)/pS396(teal) co-staining in AD CA subfields (case#22). **K** Representative PVALB(DAB)/pS396(teal) co-staining in AD DG (case#22). Scale bars: **A-C** 200 µm; **J, K** 100 µm (right), 50 µm (left).

To assess whether the regional pattern of VGAT and VGLUT1 immunoreactivity reflected a shift in the relative balance of these two presynaptic marker signals, we calculated the log2 ratio of VGLUT1 to VGAT immunoreactivity per subfield (**Fig. 5D**). In CTL, this ratio was slightly positive across most CA subfields (CA1: 0.24; CA2: 0.34; CA3: 0.26; CA4: 0.54; subiculum: 0.26), with DG showing a near-neutral value (−0.10). In AD, the ratio shifted further toward positive values in most CA subfields (CA1: 0.43; CA2: 0.22; CA3: 0.29; CA4: 0.84; subiculum: 0.17), with DG showing a negative value (−0.32). In PDD, this shift was more pronounced, with positive values in CA3, CA4, and subiculum (CA3: 0.64; CA4: 1.35; subiculum: 1.08), and a negative value in DG (−0.14) (**Fig. 5D**). As neither individual marker reached statistical significance in most of the affected subfields, this ratio-level pattern should be regarded as a secondary, composite observation rather than an independently confirmed finding.

### 3.8 Tau burden is associated with reduced inhibitory interneuron marker immunoreactivity in AD but not PDD

Spearman rank correlation between all five tau markers and five neuronal/synaptic markers (NeuN, VGAT, VGLUT1, SST, PVALB) was performed at the whole-hippocampus level within each diagnostic group separately, to determine whether tau pathology and neuronal marker density were related.

In CTL, no tau–neuronal marker correlation reached statistical significance (**Fig. 5E**). In AD hippocampus, AT8 correlated negatively with NeuN (r=−0.68, p=0.035), VGAT (r=−0.8, p=0.002), and PVALB (r=−0.86, p=0.024). GT38 showed as well a significant negative correlation with SST (r=−0.94, p=0.017) (**Fig. 5F**). At the subfield level, correlations in AD were observed in CA1 (pS422/SST, r=−0.89, p=0.033) and DG (GT38/SST, r=−0.89, p=0.033) (**Supplementary Fig. S11A-B**). These associations were illustrative of, though not directly demonstrated by, co-staining of AT8 with NeuN and PVALB, which showed predominantly AT8 single-positive neurons in regions of reduced interneuron marker immunoreactivity (**Supplementary Fig. S11C-D**). In PDD, the relationship between tau markers and neuronal density differed from AD. Rather than a negative association, pS422 showed a significant positive correlation with NeuN at the whole-hippocampus level (r=0.82, p=0.011) and in CA4 (r=0.75, p=0.025), CA3 (r=0.85, p=0.006), and CA1 (r=0.72, p=0.037). A positive correlation of pS422 with VGAT was observed in CA4 (r=0.68, p=0.05) and CA3 (r=0.72, p=0.037) as well (**Fig. 5G; Supplementary Fig. S11E-G**). These positive NeuN associations were illustrative of cellular co-staining of pS422 with NeuN and VGAT observed in PDD tissue (**Supplementary Fig. S11H, I**).

To assess whether specific interneuron populations showed differential cellular-level co-occurrence with tau pathology, we quantified the proportion of SST- and PVALB-positive interneurons co-positive for AT8 or pS396 across subfields in three AD and three PDD cases (**Fig. 5H, I**). In AD, both interneuron populations showed a greater proportion of co-positivity with pS396 than with AT8 across all subfields examined. SST/pS396 co-positivity was more prominent across CA subfields, while PVALB/pS396 co-positivity was more prominent in DG (**Fig. 5H, J, K**). This pS396 predominance was not observed in PDD (**Fig. 5I; Supplementary Fig. S11J, K**).

### 3.9 Independent transcriptomic data support reduced inhibitory neuronal markers in AD

To obtain an independent, transcript-level perspective on cell-type-associated changes in the AD hippocampus, we examined a publicly available microarray dataset (GSE48350; Affymetrix HG-U133 Plus 2.0; 43 non-demented controls, 19 AD cases). This dataset is derived from an independent cohort using bulk hippocampal tissue without subfield resolution, and cases in this dataset do not overlap with our cohort 1. It is therefore presented as external supporting context rather than as a directly linked analysis of the present cohort. Expression of the pan-neuronal marker *RBFOX3* was significantly reduced in AD relative to controls (p = 0.031; **Supplementary Fig. S12A**). All four GABAergic/interneuron transcripts examined (*SST, GAD1, GAD2, SLC32A1*) were reduced in AD relative to controls in this independent dataset (**Supplementary Fig. S12B**), whereas the excitatory neuronal transcript panel (*SLC17A7, CAMK2A, GRIN1, GRIA2*) showed a smaller and less consistent reduction (**Supplementary Fig. S12C**).

### 3.10 Distinct co-pathology profiles link tau, amyloid, and α-synuclein independent of regional identity

To assess how tau immunophenotypes are associated with amyloid-β and α-synuclein burden in AD and PDD, we examined pairwise Spearman correlations between each tau marker and 4G8 or pSyn81A across hippocampal subfields.

AT8 burden correlated positively with 4G8 (amyloid-β) burden in AD DG (r=0.66, p=0.024), CA4 (r=0.71, p=0.013), CA3 (r=0.73, p=0.009), and PHC (r=0.65, p=0.034), a pattern not observed as consistently for the other four tau markers (**Supplementary Fig. S13A-F**). AT8 and 4G8 were negatively correlated in PDD CA2 subregion (**Supplementary Fig. 13G**). Tau epitopes–pSyn81A burden correlations were largely non-significant across subfields in AD and PDD, with the exception of a positive correlation between pS396 and pSyn81A in AD CA3 (r=0.6, p=0.049) (**Supplementary Fig. S13C, H-L**).

Repeated qualitative examination of multiplex stainings showed that the frequency of tau co-pathology with α-synuclein and amyloid-β differed markedly between AD and PDD but was similar between hippocampus and PHC within each disease group. With respect to amyloid, AD hippocampus and PHC both showed AT8 and pS396 immunoreactivity enriched around and within 4G8-positive plaques, whereas pS422 and pTau217 immunoreactivity was less consistently enriched at plaque margins (**Fig. 6A; Supplementary Fig. S14A-C**). In AD, both hippocampus and PHC showed frequent co-occurrence of tau and α-synuclein pathology within individual neurons, most often as pSyn81A-immunoreactive granules within AT8-, pTau217- or pS396- positive tangles. This pattern is not observed in GT38- positive tangles. Those pSyn81A double-positive tangles can be observed as well associated to amyloid plaques (**Fig. 6B-D; Supplementary Fig. S14D-G**). In PDD, this cellular co-occurrence was less frequent in both regions, and where tau and synuclein pathology did coincide within the same neuron, it more often appeared as a Lewy body within a tau-positive tangle rather than as diffuse pSyn81A-immunoreactive granules (**Fig. 6E- H; Supplementary Fig. S14H-M**). PDD hippocampus showed minimal amyloid-β pathology by 4G8 immunoreactivity, precluding assessment of tau-amyloid spatial association in this region. PDD PHC showed low-level 4G8 immunoreactivity that was not preferentially associated with any individual tau epitope (**Fig. 6E**).

**Figure 6.**
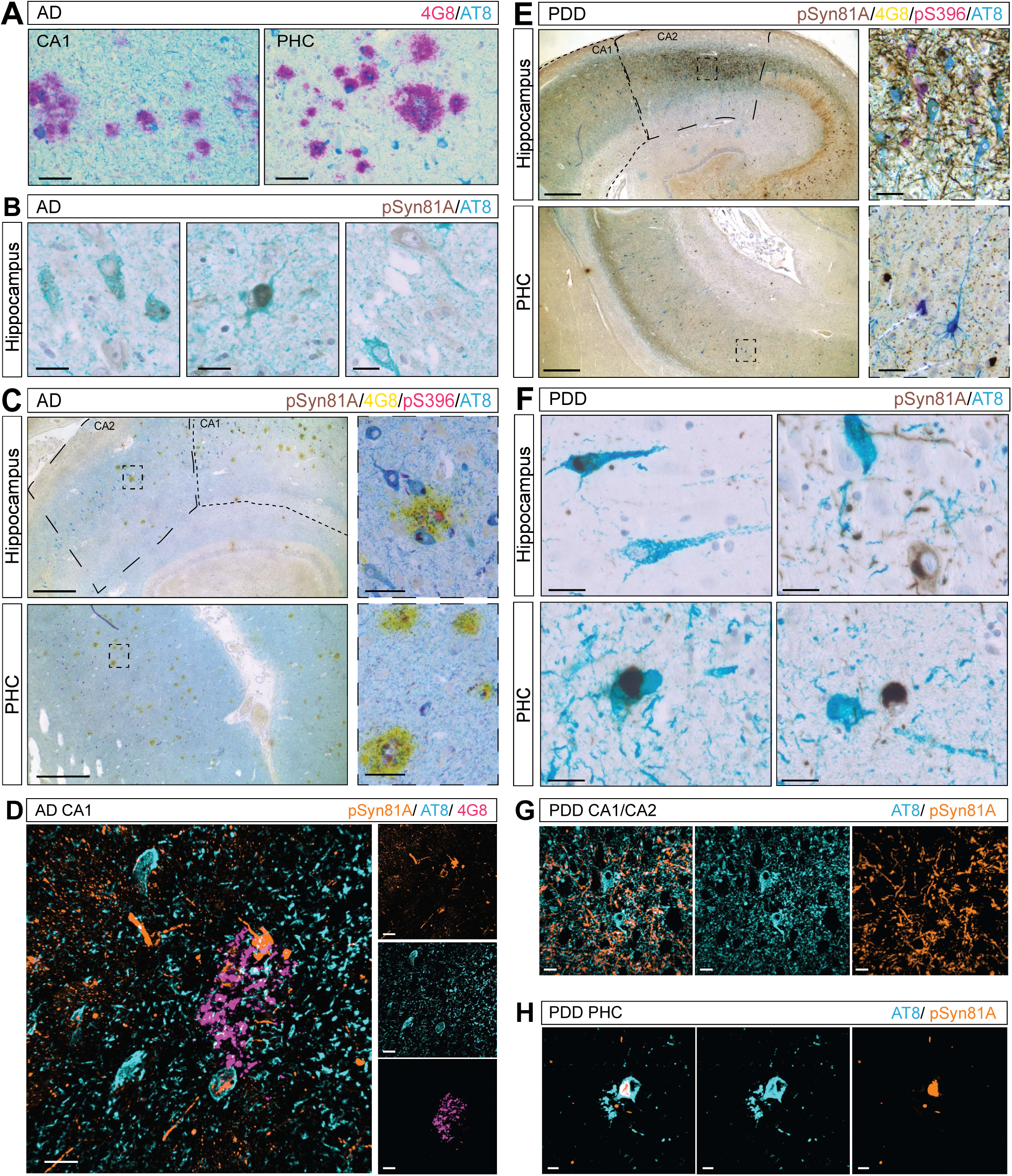
Distinct co-pathology profiles link tau, amyloid, and α-synuclein independent of regional identity. **A** Representative 4G8(purple)/AT8(teal) co-staining in AD hippocampus, illustrating AT8 immunoreactivity enriched at amyloid plaque margins (case#27). **B** Representative images of AD hippocampus, illustrating pSyn81A-immunoreactive (DAB) granules within an AT8-positive (teal) tangle, a Lewy body within an AT8- positive tangle, and spatially separate pSyn81A- and AT8-positive neurons (case #29). **C** Representative quadruplex immunohistochemistry staining of pSyn81A(DAB)/4G8(yellow)/pS396(purple)/AT8(teal) in AD hippocampus and PHC (case#27). **D** Representative immunofluorescence of pSyn81A/AT8/4G8 in AD hippocampus (case#27). **E** Representative quadruplex immunohistochemistry of pSyn81A/pS396/AT8/4G8 in PDD hippocampus and PHC (case#49). **F** Representative images of PDD, illustrating a Lewy body within an AT8-positive tangle and spatially separate pSyn81A- and AT8-positive neurons in the hippocampus and PHC (case #49). **G** Validation of AT8/pSyn81A immunofluorescence co-staining in PDD hippocampus CA1/CA2 (case#57). **H** Validation of AT8/pSyn81A immunofluorescence co-staining in PDD PHC (case#57). Scale bars: **A** 50 µm; **B**, **D**, **F** 20 µm; **C**, **E** 200 µm (right), 50 µm (left); **G**, **H** 30 µm.

## 4 Discussion

This study provides a multiscale characterisation of hippocampal and parahippocampal tau pathology across AD, PDD, DLB and non-demented CTL by integrating regional, cellular, and subcellular analysis. By combining multiplex cIHC with high-resolution confocal imaging, we identified disease-specific tangle epitope signatures and their relationships with neuronal populations and co-pathologies.

At the regional level, tau in the markers pS422, AT8, pTau217, and GT38, together with amyloid-β and α-synuclein pathologies, consistently discriminated AD from PDD and CTL, while pS396 exhibited more regional-dependent patterns. DLB showed a tau profile more similar to AD than PDD. Single-tangle immunoprofiling further revealed disease-specific heterogeneity, with AD tangles exhibiting a graded shift in phospho-tau and conformational composition from the DG to the subiculum, whereas PDD tangles were characterised by a GT38-low profile. At the cellular level, tau burden was associated with reduced inhibitory interneuron marker immunoreactivity in AD but not PDD, while confocal imaging demonstrated a conserved intra-tangle organisation in mature tangles across all disorders. Finally, local tau co-pathology with amyloid-β and α-synuclein was conserved across anatomically distinct regions within each diagnostic group despite marked differences in pathology burden.

### 4.1 Disease-specific tau signatures distinguish AD and PDD

Previous studies have shown that Alzheimer-type tau pathology occurs with variable burden across synucleinopathies, and that greater tau burden is associated with earlier dementia onset and reduced survival [27]. In contrast, α-synuclein pathology, rather than tau Braak stage, has been reported as the strongest correlate of dementia status in PDD [28]. Our findings extend this framework by showing AD and PDD differ not only in the extent of hippocampal tau pathology but also in its molecular composition. AT8, pTau217, and GT38 were consistently enriched in AD, whereas pS422 predominated in PDD and pS396 showed a more region-dependent and disease-specific distribution. PDD consistently exhibited a lower AT8 and pTau217 burden than CTL. These observations suggest that hippocampal tau pathology in PDD represents a distinct molecular profile rather than a less advanced form of AD pathology. This interpretation is consistent with emerging evidence that tau can adopt disease-specific molecular strains including distinct tau oligomer strain associated with rapidly progressive AD [29], and with a recent comparative neuropathological study identifying disease-specific tau and α-synuclein molecular signatures across AD, AD with Lewy bodies, and PDD in middle temporal gyrus and amygdala [30]. DLB exhibited a tau immunophenotype closely resembling AD, but mainly differ from PDD, consistent with imaging and fluid-biomarker evidence that DLB and PDD, despite sharing clinical features and an underlying synucleinopathy, may differ in underlying pathological burden [31].The biological basis of these distinct tau signature remains uncertain.

The regional and epitope-specific differences identified here may also have clinical implications. The relative preservation of CA subfield neuronal density in PDD, together with an AD-like tau profile in PDD PHC, is consistent with its cognitive phenotype, in which visuospatial and attentional impairment are typically more prominent than the episodic memory deficits that characterize AD [32]. In addition, the relatively low pTau217 burden observed in PDD is consistent with emerging evidence suggesting that plasma pTau217 can distinguish between AD from PDD and even between PDD and non-demented CTL in biofluid-based diagnostic settings [33].

### 4.2 Subregional gradients in tangle composition extend current models of NFT maturation

Multiplex tau epitope profiling revealed marked heterogeneity in tangle composition across disorders and hippocampal subfields. Single-tangle classification revealed a progressive shift in the relative abundance of AT8-, pS396-, and GT38-defined tangle immunophenotypes across the hippocampal subfield sequence in AD. AT8- predominant tangles were enriched in CA1 and subiculum, whereas pS396/GT38 double-positive and AT8-negative tangles enriched in the DG. This subregional gradient is consistent with current models of NFT maturation based on sequential antibody reactivity [10,34,35], in which AT8 label a broad spectrum of pretangle to mature tangle morphologies whereas GT38 is largely restricted to advanced tangle maturation morphology [11]. Our findings complement a *post-mortem* three-dimensional reconstruction of tau burden across the medial temporal lobe showing a clear anterior-to-posterior and subfield-specific gradient, centred in the transentorhinal region and CA1 [36]. Subfield-specific tangle heterogeneity is unlikely to be unique to AD. Similar regional differences in hippocampal tangle burden and morphology were reported in progressive supranuclear palsy [37], suggesting that regional differences in the capacity to support tangle formation may represent a common feature of tauopathies. Notably, many pS396-positive lacked AT8 or GT38 immunoreactivity, particularly in the DG, indicating that pS396 accumulation can occur independently of canonical AT8-defined hyperphosphorylation. In contrast, PVALB and SST neurons were more frequently associated with the pS396 immunophenotype, suggesting that tau phosphorylation trajectories differ between neuronal populations. These observations raise the possibility that distinct neuronal populations support alternative pathways of NFT maturation.

### 4.3 Intra-tangle spatial architecture as a subcellular extension of tau compositional heterogeneity

Although tau composition differed substantially between disorders, high-resolution confocal imaging demonstrated a remarkably conserved spatial organisation of phospho-tau and conformational epitopes within individual tangles. AT8 consistently occupied the outer rim, pS396 and pTau217 an intermediate core compartment, and GT38 the densely compacted core. This observations extend biochemical evidence that distinct tau phosphorylation states define biochemically and temporally discrete tau species during NFT maturation [13] by showing that mature tangles also retain a reproducible internal molecular architecture. Our observations are also consistent with single-molecule and super-resolution studies demonstrating structural heterogeneity of tau aggregates in human AD brain [38], as well as cryo-electron microscopy showing disease-specific tau filament folds across tauopathies [39,40]. Together, these findings suggest that mature NFTs are not homogeneous aggregates but highly organised molecular assemblies in which distinct phospho- or conformational epitopes retain a complex subcellular architecture.

### 4.4 Disease-specific relationship between tau signatures and neuronal vulnerability

Tau burden was associated with reduced immunoreactivity for inhibitory interneuron and presynaptic markers (SST, PVALB, VGAT) in AD, particularly for GT38 and AT8, whereas this relationship was not observed in PDD. Instead, pS422 burden in PDD showed positive associations with NeuN and VGAT, suggesting that the relationship between tau composition and neuronal integrity differs between diagnostic groups. The AD pattern is consistent with a growing literature describing parvalbumin- and somatostatin-positive GABAergic interneurons as particularly susceptible to tau pathology in the AD hippocampus [41–43]. A tau rat model further indicates differential susceptibility of PVALB and SST during circuit remodelling [42]. Recent work further suggests that tau pathology may disrupt the inhibitory circuits maintained by parvalbumin interneurons before neuronal degeneration becomes apparent, highlighting that reduced PVALB immunoreactivity may reflect functional and network-level alterations in addition to cell loss [44]. This supports the broader concept that neuronal and synaptic populations differ in their molecular response to tau pathology [45–47].

### 4.5 Co-pathology patterning reflects cell-intrinsic and disease context factors rather than network architecture

Despite marked differences in overall burden and anatomical connectivity between the hippocampus and PHC, tau pathology showed comparable spatial relationships with amyloid and α-synuclein aggregates within a given diagnostic group. These findings should be considered alongside reports supporting network-dependent, trans-synaptic tau propagation in human tauopathies. Recent work has identified oligomeric tau within synaptic pairs in human progressive supranuclear palsy [48], while computational models suggest that anatomical connectivity shape regional tau spread, with local vulnerability influencing accumulation [49]. Experimental studies similarly support interactions between pathological proteins: α-synuclein co-pathology has been reported to accelerate amyloid-driven tau accumulation in AD [50], and combined tau/amyloid/α-synuclein mouse model demonstrate synergistic enhancement on neuroinflammation and hippocampal neuron loss [51]. In addition, experimental evidence indicates bidirectional interactions between α-synuclein and tau, with α- synuclein promoting tau aggregation and hyperphosphorylation through direct cross-seeding [52,53]. Conversely, the absence of direct relationship between Lewy bodies and neuronal or synaptic loss in DLB [54], highlights the complexity of linking pathological lesions to neurodegeneration. Together these observations suggest that local co-pathology is not determined solely by regional proximity or network architecture but is shaped by cell-intrinsic or disease-context factors that in turn influence local neurodegeneration.

### 4.6 Limitations

Several limitations should be considered. The single-tangle epitope profiling, intra- tangle spatial organisation, and co-pathology stainings were performed in small case numbers (n=3 per group), with spatial organisation and co-pathology observations staying qualitative rather than formally quantified. Besides, our cross-sectional design and the limited numbers of cases used for the study preclude inference on temporal relationship or causality. Finally, IHC and IF co-positivity cannot substitute for direct biochemical or structural confirmation of the molecular identities implied by antibody reactivity.

### 4.7 Future directions

The observations reported here generate several hypotheses that can be addressed using orthogonal molecular approaches. The AD and PDD subregional and disease-specific tau signatures identified could be tested by mass spectrometry-based profiling of tau post-translational modifications, extending recent proteomic study demonstrating stage-dependent tau modification patterns across brain regions in AD [55]. The intra-tangle architecture warrants validation by higher resolution microscopy such as immuno-electron microscopy or cryo-EM. Finally, single-nucleus, spatial transcriptomic and proteomic approaches could further characterise whether the neuronal populations identified as vulnerable or resilient in this study represent distinct molecular states associated with differential tau pathology signatures [56,57].

## 5 Data availability

The data generated during the current study are available from the corresponding author upon reasonable request.

## 6 Consent for publication

All authors have consented for the publication of manuscript.

## 7 Acknowledgments

We thank the brain donors and their families for their invaluable contribution, as well as all colleagues of the National Center of Pathology Research Unit (LNS) and the teams of the Netherlands Brain Bank and the Douglas-Bell Canada Brain Bank for providing the tissue samples.

## 10 Author contributions

DSB conceived and supervised the study. SSc, MMM, FJ and MD performed the experiments. SSc performed image acquisition and processing. SSc prepared the figures. GPH and SSc performed the data analysis. SSc and DSB wrote the manuscript. All authors contributed to the final version of the manuscript.

## References

[1] DeTure MA, Dickson DW. The neuropathological diagnosis of Alzheimer’s disease. Mol Neurodegener 2019;14:32. 10.1186/s13024-019-0333-5.

[2] Hattiholi A, Hegde H, Shetty SK. Tauopathies: Emerging discoveries on tau protein, with a special focus on Alzheimer’s disease. Neuropeptides 2025;112:102536. 10.1016/j.npep.2025.102536.

[3] Biundo R, Weis L, Antonini A. Cognitive decline in Parkinson’s disease: the complex picture. Npj Parkinson’s Disease 2016;2:16018. 10.1038/npjparkd.2016.18.

[4] Agrawal S, Yu L, Leurgans SE, Nag S, Barnes LL, Bennett DA, et al. Hippocampal neuronal loss and cognitive decline in LATE-NC and ADNC among community-dwelling older persons. Alzheimers Dement 2025;21:e14500. 10.1002/alz.14500.

[5] Mak E, Su L, Williams GB, Watson R, Firbank M, Blamire A, et al. Differential Atrophy of Hippocampal Subfields: A Comparative Study of Dementia with Lewy Bodies and Alzheimer Disease. The American Journal of Geriatric Psychiatry 2016;24:136–43. 10.1016/j.jagp.2015.06.006.

[6] Yakubu AO, Atafo GI, Nwaze CE, Bakare TA, Adeniyi TO, Alare K, et al. Expanding landscape of tau pathology in neurological disorders: A narrative review guiding diagnostic development and targeted therapeutics. NeuroMarkers 2026:100167. 10.1016/j.neumar.2026.100167.

[7] J. Irwin D, I. Hurtig H. The Contribution of Tau, Amyloid-Beta and Alpha-Synuclein Pathology to Dementia in Lewy Body Disorders. J Alzheimers Dis Parkinsonism 2018;08. 10.4172/2161-0460.1000444.

[8] Drubin DG, Kirschner MW. Tau protein function in living cells. J Cell Biol 1986;103:2739–46. 10.1083/jcb.103.6.2739.

[9] Lee VM, Goedert M, Trojanowski JQ. Neurodegenerative tauopathies. Annu Rev Neurosci 2001;24:1121–59. 10.1146/annurev.neuro.24.1.1121.

[10] Moloney CM, Lowe VJ, Murray ME. Visualization of neurofibrillary tangle maturity in Alzheimer’s disease: A clinicopathologic perspective for biomarker research. Alzheimer’s & Dementia 2021;17:1554–74. 10.1002/alz.12321.

[11] Moloney CM, Rutledge MH, Labuzan SA, Peng Z, Tranovich JF, Wood AC, et al. The Neurofibrillary Tangle Maturity Scale: A Novel Framework for Tangle Pathology Evaluation in Alzheimer’s Disease 2025. 10.1101/2025.06.02.657435.

[12] Braak H, Alafuzoff I, Arzberger T, Kretzschmar H, Del Tredici K. Staging of Alzheimer disease-associated neurofibrillary pathology using paraffin sections and immunocytochemistry. Acta Neuropathol 2006;112:389–404. 10.1007/s00401-006-0127-z.

[13] Islam T, Hill E, Abrahamson EE, Servaes S, Smirnov DS, Zeng X, et al. Phospho-tau serine-262 and serine-356 as biomarkers of pre-tangle soluble tau assemblies in Alzheimer’s disease. Nat Med 2025;31:574–88. 10.1038/s41591-024-03400-0.

[14] Hely MA, Reid WGJ, Adena MA, Halliday GM, Morris JGL. The Sydney multicenter study of Parkinson’s disease: The inevitability of dementia at 20 years. Movement Disorders 2008;23:837–44. 10.1002/mds.21956.

[15] Gomperts SN. Lewy Body Dementias: Dementia With Lewy Bodies and Parkinson Disease Dementia. Continuum (Minneap Minn) 2016;22:435–63. 10.1212/CON.0000000000000309.

[16] Chu Y, Hirst WD, Federoff HJ, Harms AS, Stoessl AJ, Kordower JH. Nigrostriatal tau pathology in parkinsonism and Parkinson’s disease. Brain 2024;147:444–57. 10.1093/brain/awad388.

[17] Fixemer S, Maza MMDL, Hammer GP, Jeannelle F, Schreiner S, Gérardy J-J, et al. Microglia aggregates define distinct immune and neurodegenerative niches in Alzheimer’s disease hippocampus. 2024. 10.21203/rs.3.rs-5387511/v1.

[18] Canal-Garcia A, Branca RM, Francis PT, Ballard C, Winblad B, Lehtiö J, et al. Proteomic signatures of Alzheimer’s disease and Lewy body dementias: A comparative analysis. Alzheimer’s & Dementia 2025;21:e14375. 10.1002/alz.14375.

[19] Ma X, Zhang M, Zheng Y, Zhang X, Zhang M, Xie Y, et al. Distinct cognitive profiles differentiate dementia with lewy bodies from Alzheimer’s disease. International Psychogeriatrics 2026:100203. 10.1016/j.inpsyc.2026.100203.

[20] Compta Y, Parkkinen L, Kempster P, Selikhova M, Lashley T, Holton JL, et al. The Significance of α-Synuclein, Amyloid-β and Tau Pathologies in Parkinson’s Disease Progression and Related Dementia. Neurodegener Dis 2014;13:154–6. 10.1159/000354670.

[21] Schreiner S, Miranda de la Maza M, Darricau M, Bouvier DS. An integrated multiscale imaging workflow to resolve intracellular co-pathology in human FFPE brain tissue. Free Neuropathology 2026;7:13. 10.17879/FREENEUROPATHOLOGY-2026-9581.

[22] Kanaan NM, Cox K, Alvarez VE, Stein TD, Poncil S, McKee AC. Characterization of Early Pathological Tau Conformations and Phosphorylation in Chronic Traumatic Encephalopathy. J Neuropathol Exp Neurol 2016;75:19–34. 10.1093/jnen/nlv001.

[23] Petersen KK, Milà-Alomà M, Li Y, Du L, Xiong C, Tosun D, et al. Predicting onset of symptomatic Alzheimer’s disease with plasma p-tau217 clocks. Nat Med 2026;32:1085–94. 10.1038/s41591-026-04206-y.

[24] Montine KS, Berson E, Phongpreecha T, Huang Z, Aghaeepour N, Zou JY, et al. Understanding the molecular basis of resilience to Alzheimer’s disease. Front Neurosci 2023;17:1311157. 10.3389/fnins.2023.1311157.

[25] Bankhead P, Loughrey MB, Fernández JA, Dombrowski Y, McArt DG, Dunne PD, et al. QuPath: Open source software for digital pathology image analysis. Sci Rep 2017;7:16878. 10.1038/s41598-017-17204-5.

[26] Mai JK, Majtanik M, Paxinos G. Atlas of the human brain. 4th edition. Amsterdam Boston Heidelberg: Academic Press; 2016.

[27] Irwin DJ, Grossman M, Weintraub D, Hurtig HI, Duda JE, Xie SX, et al. Neuropathological and genetic correlates of survival and dementia onset in synucleinopathies: a retrospective analysis. Lancet Neurol 2017;16:55–65. 10.1016/S1474-4422(16)30291-5.

[28] Compta Y, Parkkinen L, O’Sullivan SS, Vandrovcova J, Holton JL, Collins C, et al. Lewy- and Alzheimer-type pathologies in Parkinson’s disease dementia: which is more important? Brain 2011;134:1493–505. 10.1093/brain/awr031.

[29] Saleem T, Möbius W, Schmitz M, Da Silva Correia A, Thomas C, Canaslan S, et al. A distinct tau oligomer strain defines the molecular and proteomic landscape of rapidly progressive Alzheimer’s disease. Acta Neuropathol 2026;151:27. 10.1007/s00401-026-02998-4.

[30] Van Der Gaag BL, Deshayes NAC, Breve JJP, Bol JGJM, Jonker AJ, Hoozemans JJM, et al. Distinct tau and alpha-synuclein molecular signatures in Alzheimer’s disease with and without Lewy bodies and Parkinson’s disease with dementia. Acta Neuropathol 2024;147:14. 10.1007/s00401-023-02657-y.

[31] Hannaway N, Zarkali A, Bhome R, Dobreva I, Thomas GEC, Veleva E, et al. Neuroimaging and plasma biomarker differences and commonalities in Lewy body dementia subtypes. Alzheimer’s & Dementia 2025;21:e70274. 10.1002/alz.70274.

[32] Oliveira HSD, Camargos ST, Cardoso F, Caramelli P. Visuospatial impairment in Parkinson’s Disease: why we should keep an eye on it. Parkinsonism & Related Disorders 2025;140:108064. 10.1016/j.parkreldis.2025.108064.

[33] Musso G, Fiorenzato E, Misenti V, Cauzzo S, Biundo R, Fogliano CA, et al. Detecting amyloid and tau pathology in Parkinson’s disease, 4R-tauopathies and control subjects with plasma pTau217. Front Neurol 2025;16:1638852. 10.3389/fneur.2025.1638852.

[34] Moloney CM, Labuzan SA, Crook JE, Siddiqui H, Castanedes-Casey M, Lachner C, et al. Phosphorylated tau sites that are elevated in Alzheimer’s disease fluid biomarkers are visualized in early neurofibrillary tangle maturity levels in the *post mortem* brain. Alzheimer’s & Dementia 2023;19:1029–40. 10.1002/alz.12749.

[35] Hamlin D, Ryall C, Turner C, Faull RLM, Murray HC, Curtis MA. Characterization of neurofibrillary tangle immunophenotype signatures to classify tangle maturity in Alzheimer’s disease. Alzheimers Dement 2024;20:4803–17. 10.1002/alz.13922.

[36] Ravikumar S, Denning AE, Lim S, Chung E, Sadeghpour N, Ittyerah R, et al. Postmortem imaging reveals patterns of medial temporal lobe vulnerability to tau pathology in Alzheimer’s disease. Nat Commun 2024;15:4803. 10.1038/s41467-024-49205-0.

[37] Best MN, Levy J, Spencer A, Min L, Olafasson E, Pinarbasi E, et al. Heterogenous Tau Pathology in the Hippocampus of Progressive Supranuclear Palsy. Alzheimer’s & Dementia 2024;20:e094106. 10.1002/alz.094106.

[38] Böken D, Cox D, Burke M, Lam JYL, Katsinelos T, Danial JSH, et al. Single-Molecule Characterization and Super-Resolution Imaging of Alzheimer’s Disease-Relevant Tau Aggregates in Human Samples. Angew Chem Int Ed 2024;63:e202317756. 10.1002/anie.202317756.

[39] Shi Y, Zhang W, Yang Y, Murzin AG, Falcon B, Kotecha A, et al. Structure-based classification of tauopathies. Nature 2021;598:359–63. 10.1038/s41586-021-03911-7.

[40] Scheres SH, Zhang W, Falcon B, Goedert M. Cryo-EM structures of tau filaments. Current Opinion in Structural Biology 2020;64:17–25. 10.1016/j.sbi.2020.05.011.

[41] Zheng J, Li H-L, Tian N, Liu F, Wang L, Yin Y, et al. Interneuron Accumulation of Phosphorylated tau Impairs Adult Hippocampal Neurogenesis by Suppressing GABAergic Transmission. Cell Stem Cell 2020;26:331–345.e6. 10.1016/j.stem.2019.12.015.

[42] Morrone CD, Lai AY, Bishay J, Hill ME, McLaurin J. Parvalbumin neuroplasticity compensates for somatostatin impairment, maintaining cognitive function in Alzheimer’s disease. Transl Neurodegener 2022;11:26. 10.1186/s40035-022-00300-6.

[43] Hernández-Frausto M, Bilash OM, Masurkar AV, Basu J. Local and long-range GABAergic circuits in hippocampal area CA1 and their link to Alzheimer’s disease. Front Neural Circuits 2023;17:1223891. 10.3389/fncir.2023.1223891.

[44] Merino-Serrais P, Plaza-Alonso S, Tapia-Gonzalez S, León-Espinosa G, DeFelipe J. Parvalbumin interneurons in the hippocampal formation of individuals with Alzheimer’s disease: a neuropathological study of abnormal phosphorylated tau in neurons. Front Neuroanat 2025;19:1571514. 10.3389/fnana.2025.1571514.

[45] Lin G, Chancellor SE, Kwon T, Woodbury ME, Doering A, Abdourahman A, et al. Cell-death pathways and tau-associated neuronal vulnerability in Alzheimer’s disease. Cell Reports 2025;44:115758. 10.1016/j.celrep.2025.115758.

[46] Torok J, Maia PD, Anand C, Raj A. Cellular underpinnings of the selective vulnerability to tauopathic insults in Alzheimer’s disease 2023. 10.1101/2023.07.06.548027.

[47] Kadamangudi S, Sanchez-Sanchez L, Kayed R, Limon A, Taglialatela G. Selective vulnerability of human synapses to soluble tau oligomers. J Alzheimers Dis 2026;110:666–84. 10.1177/13872877261416539.

[48] McGeachan RI, Keavey L, Simzer EM, Chang YY, Rose JL, Spires-Jones MP, et al. Evidence for trans-synaptic propagation of oligomeric tau in human progressive supranuclear palsy. Nat Neurosci 2025;28:1622–34. 10.1038/s41593-025-01992-5.

[49] Xiao Y, Spotorno N, An L, Bazinet V, Hansen JY, Strandberg O, et al. Brain network dynamics determine tau presence while regional vulnerability governs tau load in Alzheimer’s disease 2026. 10.21203/rs.3.rs-8687892/v1.

[50] Franzmeier N, Roemer-Cassiano SN, Bernhardt AM, Dehsarvi A, Dewenter A, Steward A, et al. Alpha synuclein co-pathology is associated with accelerated amyloid-driven tau accumulation in Alzheimer’s disease. Mol Neurodegeneration 2025;20:31. 10.1186/s13024-025-00822-3.

[51] Webster JM, Yang Y-T, Miller AT, Zane A, Scholz K, Stone WJ, et al. Tau, amyloid-β and α-synuclein co-pathologies synergistically enhance neuroinflammation and neuropathology 2024. 10.1101/2024.10.13.618101.

[52] Waxman EA, Giasson BI. Induction of Intracellular Tau Aggregation Is Promoted by α-Synuclein Seeds and Provides Novel Insights into the Hyperphosphorylation of Tau. J Neurosci 2011;31:7604–18. 10.1523/JNEUROSCI.0297-11.2011.

[53] Padilla-Godínez FJ, Vázquez-García ER, Trujillo-Villagrán MI, Soto-Rojas LO, Palomero-Rivero M, Hernández-González O, et al. α-synuclein and tau: interactions, cross-seeding, and the redefinition of synucleinopathies as complex proteinopathies. Front Neurosci 2025;19:1570553. 10.3389/fnins.2025.1570553.

[54] Hawksworth JI, Kirkby-Geddes E, Thom S, O’Neill J, Ikwue A, Wood L, et al. Lewy Bodies Are Not Associated With Neuronal or Synaptic Loss in Dementia With Lewy Bodies. Neuropathology Appl Neurobio 2026;52:e70085. 10.1111/nan.70085.

[55] Vanparys AAT, Balty C, Johanns M, Kyalu Ngoie Zola N, Herinckx G, Calsteren MV, et al. Stage-dependent tau post-translational modifications map the spatiotemporal progression of Alzheimer’s disease 2026. 10.64898/2026.04.10.717615.

[56] Dehkordi SK, Orr TC, Bodamer RH, Hudson HR, Sun V, Keene CD, et al. Spatial proteome profiling for distinguishing tangle-bearing neurons and their microenvironments from healthy neurons in Alzheimer’s disease brains. Alzheimer’s & Dementia 2024;20:e089552. 10.1002/alz.089552.

[57] Palaganas R, Sidiropoulos DN, Morris M, Stein-O’Brien GL. Mapping multipathology via spatial omic integration. Current Opinion in Biotechnology 2026;97:103398. 10.1016/j.copbio.2025.103398.

