## Supplementary Material for "Disease-specific tangle immunophenotypes distinguish hippocampal vulnerability in Alzheimer’s disease and Parkinson’s disease dementia"

Laboratoire national de santé (LNS)

1, Rue Louis Rech

L-3555 Dudelange

**Supplementary Figure S1. Aging-related tau astrogliopathy (ARTAG) detected by AT8 and pTau217 immunostaining.**

Single cIHC of serial sections stained for **A** AT8 and **B** pTau217 in CTL DG (case#5). Scale bars: **A, B** 100 µm (right), 50 µm (left).

**Supplementary Figure S2. Regional 4G8 and pSyn81A immunoreactivity in cohort 1.**

Regional staining quantification (%-DAB stained area) in dentate gyrus (DG), CA4, CA3, CA2, CA1, subiculum (SUB), parahippocampal cortex (PHC), and combined hippocampal subfields (HIPP: sum of DG, CA4, CA3, CA2, and CA1) of **A** anti-4G8 and **B** anti-pSyn81A in cohort 1. Boxplots show median and interquartile range and each point represents one case. Statistical comparisons were performed by Wilcoxon rank-sum test, with statistical significance set at $p<0.05$. Levels of significance are indicated as follows: $p<0.05$ (*), $p<0.005$ (**), and $p<0.0005$ (***), $p<0.00005$ (****).

**Supplementary Figure S3. Cohort 2 characteristics and regional tau burden across hippocampal subfields.**

**A** Demographic and neuropathological characteristics of cohort 2 (CTL n = 8, AD n=11, DLB n=14) including gender distribution, age at death, *post-mortem* delay (PMD), and Braak NFT stage. **B** Schematic table summarising tau marker morphological stainings (pretangles, mature tangles, ghost tangles, neuropil threads, neuritic plaques, tangle associated neuritic clusters (TANCs)) across pS422, AT8, pS396, pTau217, and GT38 in CTL, AD and DLB. **C** Regional AT8, 4G8, and pSyn81A immunoreactivity (%-DAB stained area) at the whole-hippocampus level in CTL, AD, and DLB. Staining quantification (%-DAB stained area) in dentate gyrus (DG), CA4, CA3, CA2, CA1, subiculum (SUB), parahippocampal cortex (PHC), and combined hippocampal subfields (HIPP: sum of DG, CA4, CA3, CA2, and CA1) of **D** anti-4G8, **E** anti-pSyn81A and **F** anti-AT8 (pS202/pT205). Each point represents one case; boxplots show median and interquartile range. Statistical comparisons were performed by Wilcoxon rank-sum test, with statistical significance set at $p < 0.05$. Levels of significance are indicated as follows: $p < 0.05$ (*), $p < 0.005$ (**), and $p < 0.0005$ (***), ****$p<0.00005$.

**Supplementary Figure S4. Antibody validation of tau markers in the hippocampus of Frontotemporal Lobar degeneration with Pick-type tau pathology (3R-tau FTLD) with Pick disease and progressive supranuclear palsy (PSP).**

Representative single cIHC staining of hippocampal subfield CA1 for **A** AT8 in 3R-tau FTLD, **B** AT8 in PSP, **C** pTau217 in 3R-tau FTLD, **D** pTau217 in PSP, **E** GT38 in 3R-tau FTLD, **F** GT38 in PSP, **G** pS396 in 3R-tau FTLD, **H** pS396 in PSP, **I** pS422 in 3R-tau FTLD, and **J** pS422 in PSP. All stainings were done n=3. Scale bars: **A- J** 100 µm.

**Supplementary Figure S5. Divergent phospho-tau and conformational immunophenotypes in cohort 2.**

Staining quantification (%-DAB stained area) in dentate gyrus (DG), CA4, CA3, CA2, CA1, subiculum (SUB), parahippocampal cortex (PHC), and combined hippocampal subfields (HIPP: sum of DG, CA4, CA3, CA2, and CA1) of **A** anti-pTau217, **B** conformation-selective tau antibody GT38, **C** anti- pS396 and **D** anti-pS422. Each point represents one case; boxplots show median and interquartile range. Statistical comparisons were performed by Wilcoxon rank-sum test, with statistical significance set at $p < 0.05$. Levels of significance are indicated as follows: $p < 0.05$ (*), $p < 0.005$ (**), and $p < 0.0005$ (***), ****$p<0.00005$.

**Supplementary Figure S6. Tau marker morphology and correlation with Braak stage across diagnostic groups.**

**A** Schematic table summarising tau marker morphological stainings (pretangles, mature tangles, ghost tangles, neuropil threads, neuritic plaques, tangle associated neuritic clusters (TANCs)) across pS422, AT8, pS396, pTau217, and GT38 in CTL, AD and PDD. **B-F** Spearman correlation of tau burden versus Braak NFT stage separated by diagnostic group for **B** anti-pS422, **C** anti-AT8, **D** anti-pS396, **E** anti-pTau217 and **F** anti-GT38.

**Supplementary Figure S7. Pairwise correlation of tau markers across hippocampal subfields in AD and PDD.**

**A–F** Pairwise tau marker correlation matrices of AD hippocampal subfields and the PHC. **G-L** Pairwise tau marker correlation matrix of PDD hippocampal subfields and the PHC. Each cell reports the Spearman correlation coefficient (ρ) between %DAB-stained area for the two indicated tau markers, calculated across all cases within the diagnostic group. Colour scale indicates the direction and magnitude of ρ (blue= positive, red= negative, range 1 to -1). Correlations were considered significant at $p<0.05$ (two-sided). Non-significant correlations are marked with a grey dot. *P* values were obtained from two-sided Spearman rank correlation tests performed separately for each pair of markers.

**Supplementary Figure S8. Sequential distribution and co-localisation of tau epitopes in control and AD hippocampus**

**A** Representative triplex cIHC of pS422(red)/pS396(teal)/GT38(green) in CTL CA1 (case#9). **B** Representative triplex chromogenic cIHC of pS422(red)/pS396(teal)/pTau217(green) in CTL CA1 (case#9). **C** Representative triplex cIHC of pTau217(red)/pS396(teal)/AT8(green) in CTL CA1 (case#9). **D** Duplex cIHC staining for AT8 (purple) and GFAP (green) (case#5). **E** Duplex cIHC staining for pTau217 (purple) and GFAP (green) (case#5). **F** Triplex IF staining of pTau217 (orange), AT8 (magenta) and GFAP (green) demonstrating intracellular tau pathology in astrocytes (case#5). **G**, **H** Representative duplex cIHC of **G** pTau217(purple)/AT8(teal) and **H** pTau217(purple)/GT38(teal) in AD CA1 (case#32). Scale bars: **A-C** 100 µm (right), 50 µm (left); **D, E** 100 µm (right), 50 µm (left); **F** 20 µm; **G**, **H 50** µm.

**Supplementary Figure S9. Intra-tangle organisation of tau species in DLB and AD revealed by multiplex immunohistochemistry and confocal imaging.**

**A** Representative multiplex cIHC stainings of GT38(DAB)/ pS396(purple)/AT8(teal) illustrating single-tangle immunophenotypes in DLB hippocampus (case#70). **B** Representative triplex IF of pTau217(purple)/AT8(teal)/ pS396(orange) in AD CA1, supporting the intra-tangle organisation shown in Fig. 4H (case#32). **C** High-resolution confocal z-stack imaging of GT38(orange)/ pS396(purple)/AT8(teal) triplex-stained DLB CA1 showing subcellular NFT organisation: AT8 at the outer rim, pS396 in an inner compartment, GT38 innermost (case#70). Scale bars: **A** 50 µm; **B** 30 µm; **C** 30 µm (overview), 15 µm (inset).

**Supplementary Figure S10. Interneuron and excitatory neuron markers are altered in AD and PDD hippocampus.**

Staining quantification (%-DAB stained area) in dentate gyrus (DG), CA4, CA3, CA2, CA1, subiculum (SUB), parahippocampal cortex (PHC), and combined hippocampal subfields (HIPP: sum of DG, CA4, CA3, CA2, and CA1) of **A** SST and **B** PVALB in CTL, AD and PDD. Boxplots show median and interquartile range and each point represents one case. Statistical comparisons were performed by Wilcoxon rank-sum test, with statistical significance set at $p<0.05$. Levels of significance are indicated as follows: $p<0.05$ (*), $p<0.005$ (**), and $p<0.0005$ (***), $p<0.00005$ (****).

**Supplementary Figure S11. Tau pathology correlates with neuronal loss across hippocampal subfields.**

**A, B** Subfield-level correlation between tau burden (%DAB-stained area) and neuronal density in AD CA1 and DG. Each cell reports the Spearman correlation coefficient (ρ) between %DAB-stained area for the two indicated tau markers, calculated across all cases within the diagnostic group. Colour scale indicates the direction and magnitude of ρ (blue= positive, red= negative, range 1 to -1). Correlations were considered significant at $p<0.05$ (two-sided). Non-significant correlations are marked with a grey dot. *P* values were obtained from two-sided Spearman rank correlation tests performed separately for each pair of markers. **C** Representative AT8(purple)/NeuN (teal) co-staining in AD CA1 (case#28). **D** Representative AT8(yellow)/PVALB(teal) co-staining in AD CA1 (case#27). **E–G** Subfield-level tau–neuron correlation of PDD CA4, CA3 and CA1. Each cell reports the Spearman correlation coefficient (ρ) between %DAB-stained area for the two indicated tau markers, calculated across all cases within the diagnostic group. Colour scale indicates the direction and magnitude of ρ (blue= positive, red= negative, range 1 to -1). Correlations were considered significant at $p<0.05$ (two-sided). Non-significant correlations are marked with a grey dot. *P* values were obtained from two-sided Spearman rank correlation tests performed separately for each pair of markers. **H, I** Representative pS422(red)/NeuN(teal) and pS422(red)/VGAT(teal) multiplex cIHC in PDD CA1 (case#57). **J, K** Representative SST(DAB)/PVALB(DAB)–pS396(teal) cIHC in PDD

CA1 (case#57). Scale bars: **C** 100 µm (right), 50 µm (left); **D** 100 µm (right), 30 µm (left); **H-K** 50 µm.

**Supplementary Figure S12. Independent replication of neuronal and interneuron marker changes in Alzheimer's disease using GSE48350.**

**A** *RBFOX3* expression in AD vs. control, GSE48350 (independent cohort). **B** GABAergic/interneuron transcript panel (*SST, GAD1, GAD2, SLC32A1*) in AD vs. control, GSE48350 (independent cohort). **C** Excitatory neuronal transcript panel (*SLC17A7, CAMK2A, GRIN1, GRIA2*) in AD vs. control, GSE48350 (independent cohort). Group comparisons were performed using two-sided Welch's t-test (unpaired, unequal variances). Significance is indicated as $p<0.05$ (*), $p<0.005$ (**), $p<0.0005$ (***), $p<0.00005$ (****).

**Supplementary Figure S13. Subfield-level correlation between tau pathology and co-pathology burden.**

**A–F** Subfield-level tau (pS422, AT8, pS396, GT38, pTau217) and co-pathology (4G8, pSyn81A) correlation in AD. **G-L** Subfield-level tau (pS422, AT8, pS396, GT38, pTau217) and co-pathology (4G8, pSyn81A) correlation in PDD. Each cell reports the Spearman correlation coefficient (ρ) between %DAB-stained area for the two indicated tau markers, calculated across all cases within the diagnostic group. Colour scale indicates the direction and magnitude of ρ (blue= positive, red= negative, range 1 to -1). Correlations were considered significant at $p<0.05$ (two-sided). Non-significant correlations are marked with a grey dot. *P* values were obtained from two-sided Spearman rank correlation tests performed separately for each pair of markers.

**Supplementary Figure S14. Spatial relationships between tau, α-synuclein and amyloid-β pathology.**

**A–C** Representative images of AD hippocampus showing tau/4G8 plaque margin enrichment (case#24, #27, #28). **D-G** Representative images of AD showing tau/pSyn81A granule co-occurrence within different tau-stained tangles (case#29). **H–M** Representative images of PDD hippocampus and PHC, showing different tau/α-synuclein cellular relationship, including Lewy body within tangle morphology (case#47). Scale bars: **A-M** 50 μm.

### Supplementary Tables

***Supplementary Table 1: Cohort demographics and neuropathological characteristics.***
*Post-mortem brain tissue cases included in this study with anonymised case identifiers, neuropathological diagnosis (CTL, control; AD, Alzheimer's disease; PDD, Parkinson's disease dementia; DLB, dementia with Lewy bodies; FTD, Frontotemporal dementia; PSP, Progressive supranuclear palsy), sex, age at death (years), post-mortem delay (PMD, hours), neuropathological scores including ABC score and Braak neurofibrillary tangle stage, and additional clinical annotations (AGD, argyrophilic grain disease; PART, primary age-related tauopathy; ARTAG, age-related tau astrogliopathy; LATE, limbic-predominant age-related TDP-43 encephalopathy). Brain bank of origin is indicated (Cohort 1, Netherlands Brain Bank; Cohort 2, Montreal). N/A, not annotated.*

| CASE | PATHOLOGICAL DIAGNOSIS | SEX | AGE AT DEATH | PMD (hours) | Cognitive scoring | SCORE | Clinical Info | BRAIN BANK |
|---|---|---|---|---|---|---|---|---|
| #1 | CTL | F | 87 | 6.25 | MMSE 30/30 | A2B1C1, Braak 2 | AGD | Netherland (cohort 1) |
| #2 | CTL | F | 102 | 3.55 | MMSE 30/30 | A1B2C0, Braak 3 | PART, ARTAG | Netherland (cohort 1) |
| #3 | CTL | F | 79 | 6.0 | N/A | A2B2C2, Braak 3 | ARTAG | Netherland (cohort 1) |
| #4 | CTL | M | 96 | 4.5 | N/A | A3B2C2, Braak 3 | ARTAG | Netherland (cohort 1) |
| #5 | CTL | M | 87 | 6.25 | N/A | A1B2C0, Braak 3 | AGD, ARTAG | Netherland (cohort 1) |
| #6 | CTL | F | 104 | 7.33 | N/A | A2B2C1, Braak 4 | ARTAG | Netherland (cohort 1) |
| #7 | CTL | M | 91 | 6.35 | N/A | A1B1C0, Braak 2 | ARTAG | Netherland (cohort 1) |
| #8 | CTL | F | 92 | 6.0 | N/A | A3B2C2, Braak 3 | AGD, ARTAG | Netherland (cohort 1) |
| #9 | CTL | F | 90 | 7.3 | N/A | A1B2C0, Braak 3 | ARTAG | Netherland (cohort 1) |
| #10 | CTL | M | 89 | 32.25 | N/A | A0B0C0 | N/A | Montreal (cohort 2) |
| #11 | CTL | F | 83 | 35.75 | N/A | A0B0C0 | N/A | Montreal (cohort 2) |
| #12 | CTL | M | 72 | 7.48 | N/A | A0B0C0 | N/A | Montreal (cohort 2) |
| #13 | CTL | M | 80 | 13 | N/A | A1B1C1 | N/A | Montreal (cohort 2) |
| #14 | CTL | M | 68 | 23.37 | N/A | A0B1C0 | N/A | Montreal (cohort 2) |
| #15 | CTL | M | 83 | 16.83 | N/A | A0B0C0 | N/A | Montreal (cohort 2) |
| #16 | CTL | F | 86 | 5.75 | N/A | A1B1C1 | N/A | Montreal (cohort 2) |
| #17 | CTL | M | 76 | 19.75 | N/A | A0B0c0 | N/A | Montreal (cohort 2) |
| #18 | CTL | F | 89 | 23.58 | N/A | A1B1C0 | N/A | Montreal (cohort 2) |
| #20 | AD | F | 86 | 3.5 | N/A | A3B3C3, Braak 5 | ARTAG, LATE | Netherland (cohort 1) |
| #21 | AD | F | 98 | 6.5 | N/A | A2B2C2, Braak 4 | AGD | Netherland (cohort 1) |
| #22 | AD | M | 38 | 5.45 | N/A | A3B3C3, Braak 6 | N/A | Netherland (cohort 1) |
| #23 | AD | M | 64 | 4.58 | N/A | A3B3C3, Braak 5 | N/A | Netherland (cohort 1) |
| #24 | AD | F | 70 | 7.25 | N/A | A3B3C3, Braak 6 | LATE | Netherland (cohort 1) |
| #25 | AD | F | 53 | 6.3 | N/A | A3B3C3, Braak 6 | N/A | Netherland (cohort 1) |
| #26 | AD | M | 74 | 5.25 | N/A | A3B3C3, Braak 6 | ARTAG | Netherland (cohort 1) |
| #27 | AD | F | 86 | 6.0 | N/A | A3B3C3, Braak 5 | ARTAG | Netherland (cohort 1) |

| | | | | | | | | |
|---|---|---|---|---|---|---|---|---|
| #28 | AD | F | 90 | 5.55 | N/A | A3B3C2, Braak 5 | ARTAG | Netherland (cohort 1) |
| #29 | AD | M | 82 | 8.35 | MMSE 16/30 | A3B3C3, Braak 6 | AGD, LATE, ARTAG | Netherland (cohort 1) |
| #30 | AD | M | 79 | 4.1 | MMSE 17/30 | A3B2C1, Braak 4 | ARTAG, LATE | Netherland (cohort 1) |
| #31 | AD | M | 78 | 10.2 | MMSE 29/30 | A3B2C2 Braak 4 | ARTAG | Netherland (cohort 1) |
| #32 | AD | F | 65 | 8.25 | MoCA 21/30 | A3B3C3, Braak 5 | ARTAG | Netherland (cohort 1) |
| #33 | AD | M | 90 | 26.12 | N/A | A3B2C3 | N/A | Montreal (cohort 2) |
| #34 | AD | M | 82 | 11.5 | N/A | A2B2C2 | N/A | Montreal (cohort 2) |
| #35 | AD | M | 93 | 13.87 | N/A | A2B3C2 | N/A | Montreal (cohort 2) |
| #36 | AD | F | 81 | 23.75 | N/A | A2B3C2 | N/A | Montreal (cohort 2) |
| #37 | AD | M | 95 | 29.5 | N/A | A2B3C2 | N/A | Montreal (cohort 2) |
| #38 | AD | M | 77 | 19.42 | N/A | A2B3C2 | N/A | Montreal (cohort 2) |
| #39 | AD | M | 90 | 14.6 | N/A | A2B2C2 | N/A | Montreal (cohort 2) |
| #40 | AD | M | 90 | 31.98 | N/A | A2B2C3 | N/A | Montreal (cohort 2) |
| #41 | AD | F | 87 | 17 | N/A | A1B2C2 | N/A | Montreal (cohort 2) |
| #42 | AD | F | 83 | 25 | N/A | A2B3C2 | N/A | Montreal (cohort 2) |
| #43 | AD | M | 91 | 25 | N/A | A2B3C2 | N/A | Montreal (cohort 2) |
| #44 | AD | M | 87 | 36.75 | N/A | A2B3C2 | N/A | Montreal (cohort 2) |
| #45 | AD | M | 87 | 10.8 | N/A | A2B3C2 | N/A | Montreal (cohort 2) |
| #46 | AD | F | 96 | 26.5 | N/A | A2B3C2 | N/A | Montreal (cohort 2) |
| #47 | PDD | M | 76 | 5.15 | N/A | Braak 1 | N/A | Netherland (cohort 1) |
| #48 | PDD | F | 75 | 6.10 | N/A | Braak 1 | N/A | Netherland (cohort 1) |
| #49 | PDD | M | 71 | 4.35 | N/A | Braak 2 | N/A | Netherland (cohort 1) |
| #50 | PDD | M | 73 | 5.35 | N/A | Braak 1 | N/A | Netherland (cohort 1) |
| #51 | PDD | M | 72 | 4.0 | N/A | Braak 1 | N/A | Netherland (cohort 1) |
| #52 | PDD | F | 71 | 9.05 | N/A | A0B1C0, Braak 2 | N/A | Netherland (cohort 1) |
| #53 | PDD | M | 85 | 5.15 | N/A | A1B2C1 Braak 1 | ARTAG | Netherland (cohort 1) |
| #54 | PDD | F | 73 | 6.1 | N/A | A1B1C0, Braak 2 | ARTAG | Netherland (cohort 1) |
| #55 | PDD | F | 88 | 6.05 | N/A | A1B1C0, Braak 2 | ARTAG | Netherland (cohort 1) |
| #56 | PDD | F | 69 | 7.05 | N/A | A1B1C0, Braak 1 | N/A | Netherland (cohort 1) |
| #57 | PDD | M | 72 | 4.0 | N/A | A0B1C0, Braak 1 | ARTAG | Netherland (cohort 1) |
| #58 | PDD | F | 66 | 5.5 | MMSE 23/30 | A2B0C0, Braak 0 | N/A | Netherland (cohort 1) |

| | | | | | | | | |
|---|---|---|---|---|---|---|---|---|
| #59 | PDD | M | 79 | 6.2 | N/A | A1B1C0, Braak 0 | PART, ARTAG, AGD | Netherland (cohort 1) |
| #60 | PDD | F | 83 | 6.05 | N/A | A0B1C0, Braak 1 | N/A | Netherland (cohort 1) |
| #61 | PDD | F | 80 | 6.1 | N/A | A1B2C0, Braak 2 | N/A | Netherland (cohort 1) |
| #62 | DLB | M | 70 | 41.33 | N/A | N/A | N/A | Montreal (cohort 2) |
| #63 | DLB | M | 75 | 21 | N/A | N/A | N/A | Montreal (cohort 2) |
| #64 | DLB | F | 76 | 29.25 | N/A | N/A | N/A | Montreal (cohort 2) |
| #65 | DLB | M | 64 | 17.25 | N/A | N/A | N/A | Montreal (cohort 2) |
| #66 | DLB | M | 76 | 29.25 | N/A | N/A | N/A | Montreal (cohort 2) |
| #67 | DLB | F | 83 | 17.75 | N/A | N/A | N/A | Montreal (cohort 2) |
| #68 | DLB | M | 89 | 24.5 | N/A | N/A | N/A | Montreal (cohort 2) |
| #69 | DLB | M | 83 | 6.25 | N/A | N/A | N/A | Montreal (cohort 2) |
| #70 | DLB | F | 88 | 26 | N/A | N/A | N/A | Montreal (cohort 2) |
| #71 | DLB | M | 82 | 22 | N/A | N/A | N/A | Montreal (cohort 2) |
| #72 | 3R-tau FTLD | M | 72 | 5.35 | N/A | A1B1C0 | N/A | Netherland |
| #73 | FTD Pick's disease | F | 74 | 9.05 | N/A | A1B1C0, Braak 1 | N/A | Netherland |
| #74 | FTD Pick's disease | F | 72 | 9.1 | MMSE 12/30 | A1B0C0, Braak 0 | N/A | Netherland |
| #75 | PSP | F | 72 | 5.45 | N/A | A1B1C0, Braak 2 | N/A | Netherland |
| #76 | PSP | M | 71 | 4.45 | N/A | A0B1C0, Braak 1 | ARTAG | Netherland |
| #77 | PSP | F | 76 | 5.2 | N/A | A3B2C1, Braak 3 | ARTAG | Netherland |

***Supplementary Table 2: Primary antibodies, secondary antibodies and compounds used in this study.*** *Antibodies and compounds used for immunohistochemical and immunofluorescence staining, including antibody name, host species, source, catalogue number (Cat#) and Research Resource Identifier (RRID), and dilution applied for DAKO Omnis (D) and UltraDiscovery (UD). N/A, not annotated.*

| ANTIBODIES | SOURCE | IDENTIFIER | DILUTION |
|---|---|---|---|
| PRIMARY ANTIBODIES | | | |
| Mouse monoclonal anti-β-Amyloid, clone 4G8 | BioLegend | Cat# 800712, RRID: AB_2734548 | 1:500, Low pH (D) 1:500 (UD) |
| Mouse monoclonal anti-GT38 | Abcam | Cat# 760–4345, RRID: N/A | 1:200, High pH (D) 1:500 (UD) |
| Mouse monoclonal anti-NeuN, clone A60 | Millipore | Cat# MAB377, RRID: AB_2298772 | 1:200 (UD) |
| Rabbit polyclonal anti-PVALB | Atlas Antibodies | Cat# HPA048536, RRID: AB_2680433 | 1:200 (UD) |
| Mouse monoclonal anti-phospho-α-Synuclein (Ser129), clone 81A | Millipore | Cat# MABN826, RRID:AB_2904158 | 1:200, Low pH (D) 1:500 (UD) |
| Mouse monoclonal anti- phospho-Tau (Ser202, Thr205) (AT8) | Thermo Fisher Scientific | Cat# MN1020, RRID: AB_223647 | 1:400, Low pH (D) 1:1000 (UD) |
| Rabbit polyclonal anti-phospho-Tau (Thr217) | Innovative Research | Cat# 44-744, RRID: AB_1502121 | 1:500 (UD) |
| Rabbit polyclonal anti-phospho-Tau (Ser396) | Innovative Research | Cat# 44-752G, RRID: AB_1502108 | 1:1000, Low pH (D) 1:1000 (UD) |
| Rabbit polyclonal anti-phospho-Tau (Ser422) | Innovative Research | Cat# 44-764G, RRID: AB_1502115 | 1:500, Low pH (D) 1:200 (UD) |
| Rabbit polyclonal anti-SST | Sigma-Aldrich | Cat# HPA019472, RRID: AB_1857360 | 1:200 (UD) |
| Rabbit polyclonal anti-VGAT | Sigma-Aldrich | Cat# HPA058859, RRID: AB_2683836 | 1:200 (UD) |
| Rabbit polyclonal anti-VGLUT1 | Sigma-Aldrich | Cat# HPA063679, RRID: AB_2685086 | 1:200 (UD) |
| SECONDARY ANTIBODIES | | | |
| DISCOVERY CM DAB kit | Roche | *Cat# 760-159, RRID: N/A* | Ready to use* |
| DISCOVERY Purple Kit | Roche | *Cat# 760-229, RRID: N/A* | Ready to use* |
| DISCOVERY Teal HRP kit | Roche | *Cat# 760-247, RRID: N/A* | Ready to use* |
| DISCOVERY Yellow Kit | Roche | *Cat# 760-239, RRID: N/A* | Ready to use* |
| DISCOVERY Green HRP Kit | Roche | *Cat# 760-271, RRID: N/A* | Ready to use* |
| DISCOVERY RED Kit | Roche | *Cat# 760-228, RRID: N/A* | Ready to use* |
| DISCOVERY Rhodamine 6G Kit | Roche | *Cat# 760-233, RRID: N/A* | Ready to use* |
| DISCOVERY Cy5 Kit | Roche | *Cat# 760-238, RRID: N/A* | Ready to use* |
| DISCOVERY FAM Kit | Roche | *Cat# 760-243, RRID: N/A* | Ready to use* |
| DISCOVERY QD DAPI (RUO) | Roche | *Cat # 760-4196, RRID: N/A* | Ready to use* |
| COMPOUNDS | | | |
| Reaction buffer concentrate (10X) | Roche | Cat# 950–300, *RRID : N/A* | Ready to use* |

| | | | |
|---|---|---|---|
| Haematoxylin II | Roche | *Cat# 790–2208,*<br>*RRID : N/A* | Ready to use* |
| Bluing reagent | Roche | *Cat# 760–2039,*<br>*RRID : N/A* | Ready to use* |
| 10X EZ Prep solution | Roche | *Cat# 950–102,*<br>*RRID : N/A* | Ready to use* |
| DISCOVERY Cell Conditioning 1 (CC1) | Roche | *Cat# 950–224,*<br>*RRID : N/A* | Ready to use* |
| DISCOVERY Cell Conditioning 2 (CC2) | Roche | *Cat# 950–223,*<br>*RRID : N/A* | Ready to use* |
| DISCOVERY Inhibitor | Roche | *Cat# 950–300,*<br>*RRID : N/A* | Ready to use* |
| EnVision™FLEX Target Retrieval Solution high pH buffer | Agilent Dako | *Cat# K8004,*<br>*RRID : N/A* | Ready to use* |
| EnVision™FLEX Target Retrieval Solution low pH buffer | Agilent Dako | *Cat# K8005,*<br>*RRID : N/A* | Ready to use* |
| EnVision™FLEX Antibody Diluent | Agilent Dako | *Cat# K8006,*<br>*RRID : N/A* | Ready to use* |
| EnVision™FLEX Detection kit | Agilent Dako | *Cat# K8000,*<br>*RRID : N/A* | Ready to use* |
| Haematoxylin | Agilent Dako | *Cat# GC808,*<br>*RRID : N/A* | Ready to use* |

*Vials ready-to-use

***Supplementary Table 3: HALO image analysis software threshold settings.***
*Parameters used to define tissue segmentation and staining thresholds for quantitative area fraction analysis in HALO software (Indica Labs). HTX, haematoxylin counterstain channel used for tissue detection; OD, optical density threshold applied to define DAB-positive immunoreactivity. Settings are reported per marker and staining run to ensure reproducibility.*

| Antibody | HTX STAIN COLOR | HTX Min OD | DAB STAIN COLOR | DAB Min OD | BACKGROUND STAIN COLOR | MINIMUM TISSUE OD |
|---|---|---|---|---|---|---|
| 4G8 | 0.097;0.097; 0.066 | 0.078;0.149 6; 0.3165 | 0.097;0.097; 0.066 | 0.15;0.275; 0.5054 | 0.058 ;0.066; 0.054 | 0.037 |
| pSyn81A | 0.097;0.097; 0.066 | 0.078;0.149 6; 0.3165 | 0.097;0.097; 0.066 | 0.15;0.275; 0.55 | 0.058 ;0.066; 0.054 | 0.037 |
| AT8 | 0.097;0.097; 0.066 | 0.078;0.149 6; 0.3165 | 0.200;0.342; 0.402 | 0.15 ;0.275; 0.55 | 0.058 ;0.066; 0.054 | 0.037 |
| pS422 | 0.097;0.097; 0.066 | 0.078;0.149 6; 0.3165 | 0.200; 0.342; 0.402 | 0.15;0.275; 0.55 | 0.058 ;0.066; 0.054 | 0.037 |
| pS396 | 0.097;0.097; 0.066 | 0.078;0.149 6; 0.3165 | 0.200; 0.342; 0.402 | 0.15;0.275; 0.55 | 0.058 ;0.066; 0.054 | 0.037 |
| pTau217 | 0.097;0.097; 0.066 | 0.078;0.149 6; 0.3165 | 0.142; 0.276; 0.289 | 0.025;0.27 5; 0.55 | 0.058 ;0.066; 0.054 | 0.037 |
| GT38 | 0.097;0.097; 0.066 | 0.078;0.149 6; 0.3165 | 0.200; 0.342; 0.402 | 0.15;0.275; 0.55 | 0.058 ;0.066; 0.054 | 0.037 |
| NeuN | 0.097;0.097; 0.066 | 0.078;0.149 6; 0.3165 | 0.123;0.168; 0.054 | 0.15;0.275; 0.55 | 0.058 ;0.066; 0.054 | 0.037 |
| SST | 0.097;0.097; 0.066 | 0.078;0.149 6; 0.3165 | 0.097;0.097; 0.066 | 0.03;0.275; 0.55 | 0.058 ;0.066; 0.054 | 0.037 |
| PVALB | 0.097;0.097; 0.066 | 0.078;0.149 6; 0.3165 | 0.097;0.097; 0.066 | 0.001;0.27 5;0.55 | 0.058 ;0.066; 0.054 | 0.037 |
| VGAT | 0.097;0.097; 0.066 | 0.078;0.149 6; 0.3165 | 0.086; 0.130; 0.066 | 0.04;0.610; 1.242 | 0.058 ;0.066; 0.054 | 0.037 |
| VGLUT1 | 0.097;0.097; 0.066 | 0.078;0.149 6; 0.3165 | 0.086; 0.130; 0.066 | 0.09; 0.610; 1.242 | 0.058 ;0.066; 0.054 | 0.037 |

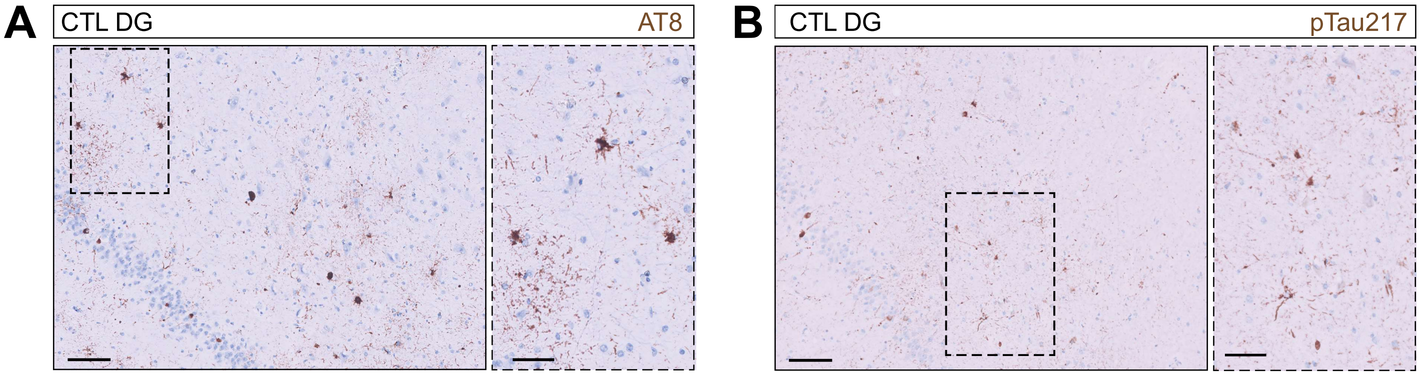


**Supplementary Figure S1**

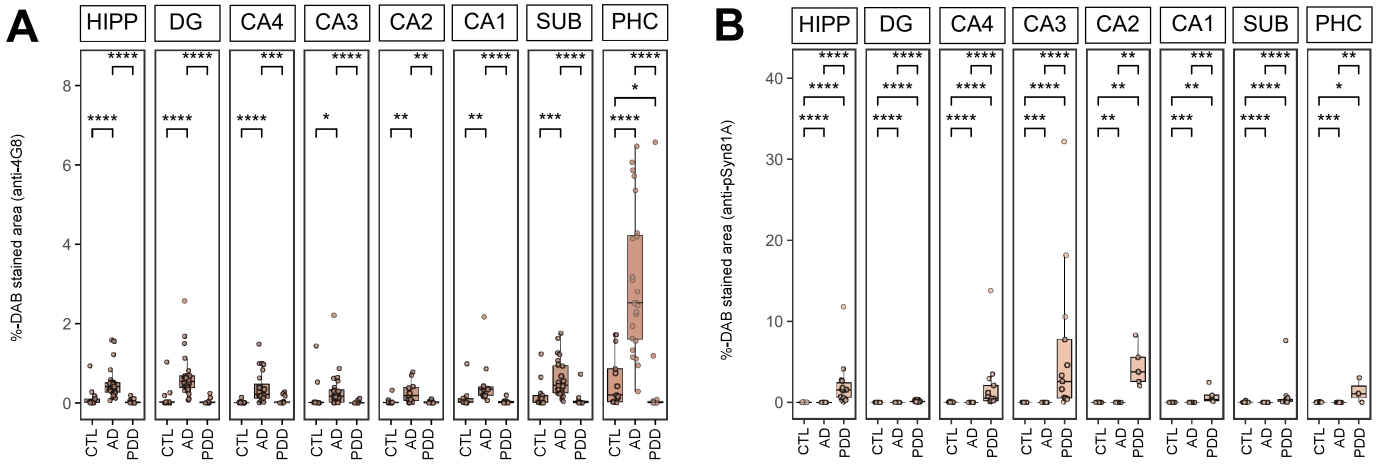


**Supplementary Figure S2**

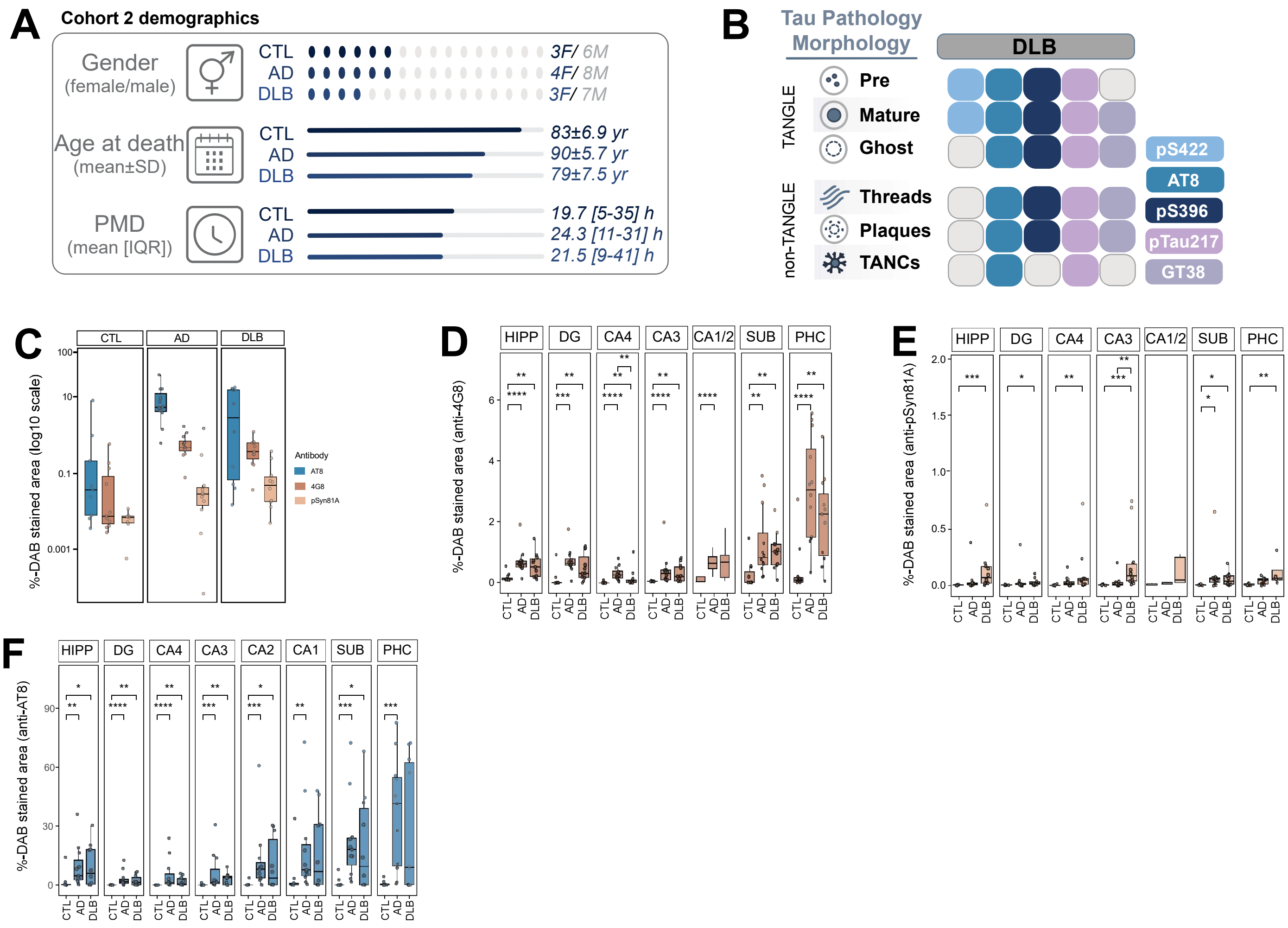


**Supplementary Figure S3**

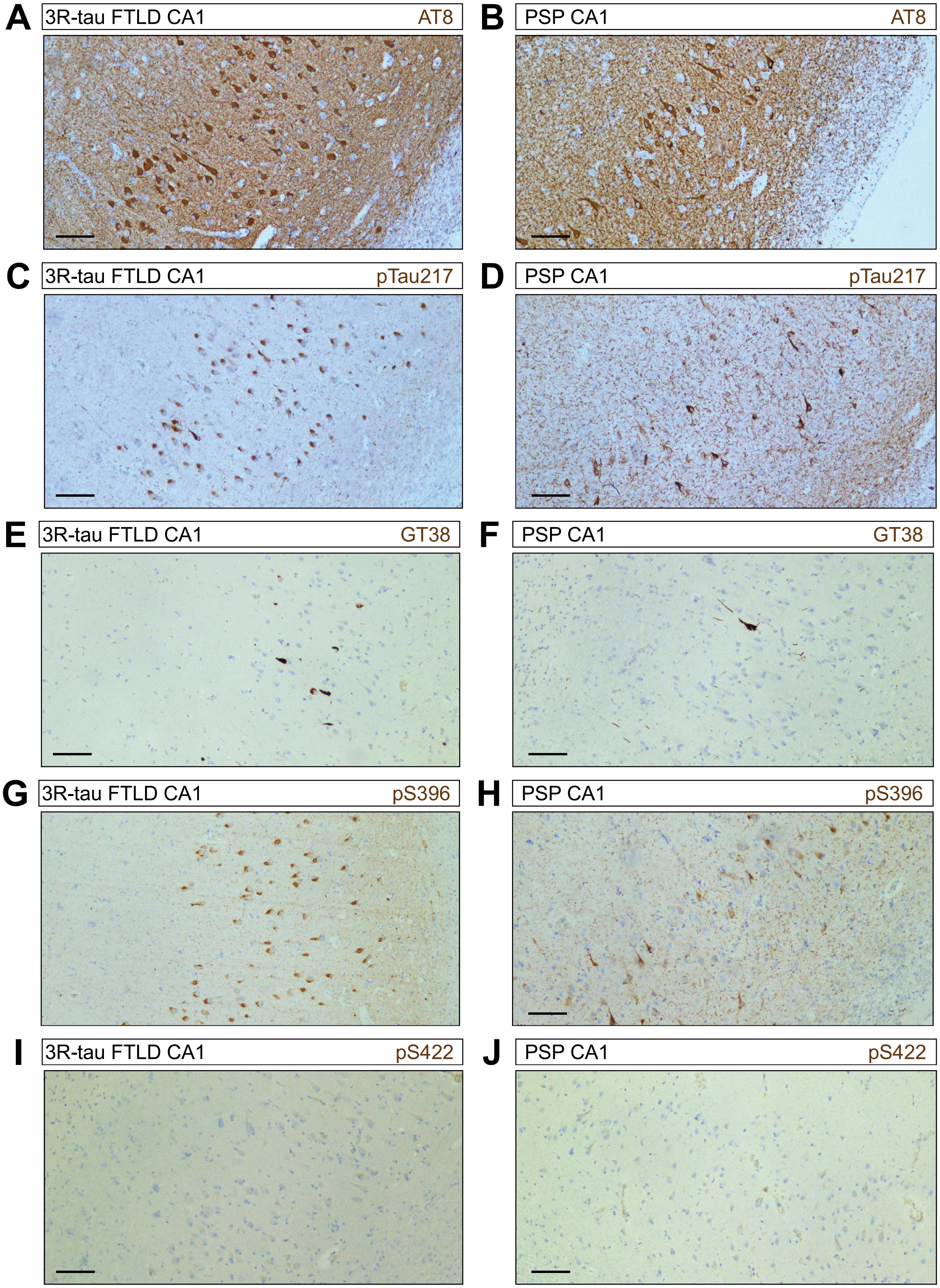


**Supplementary Figure S4**

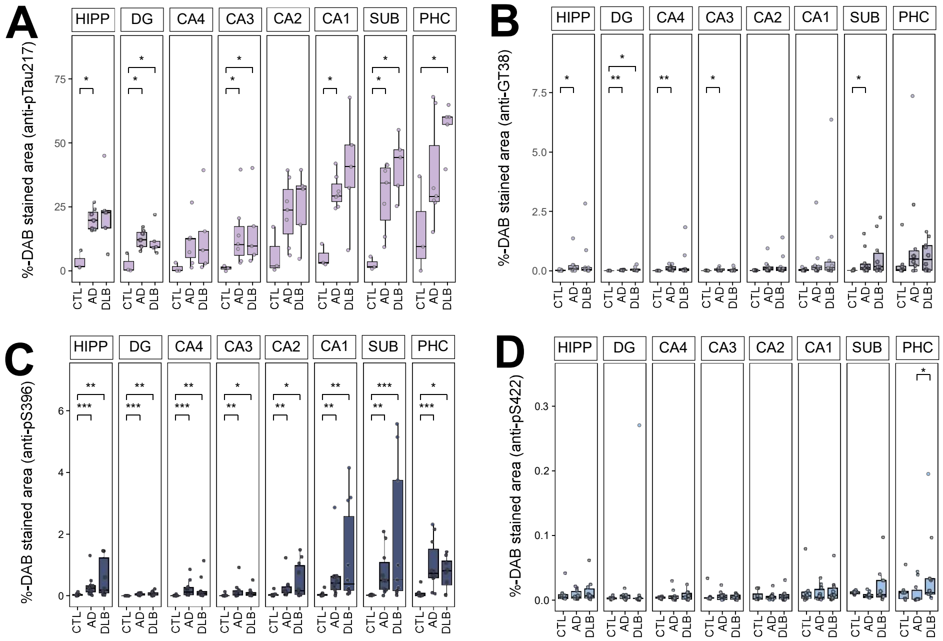


**Supplementary Figure S5**

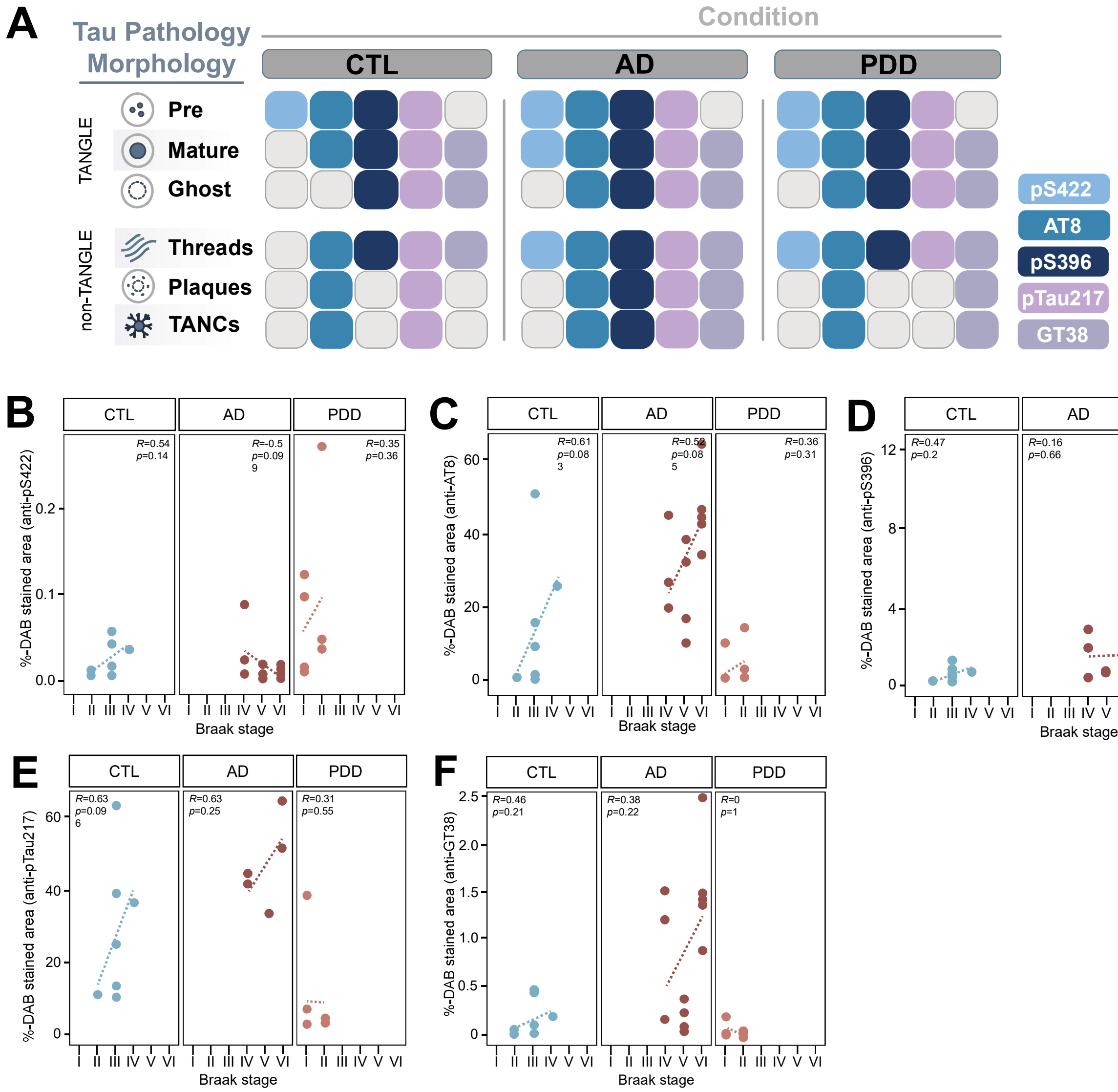


**Supplementary Figure S6**

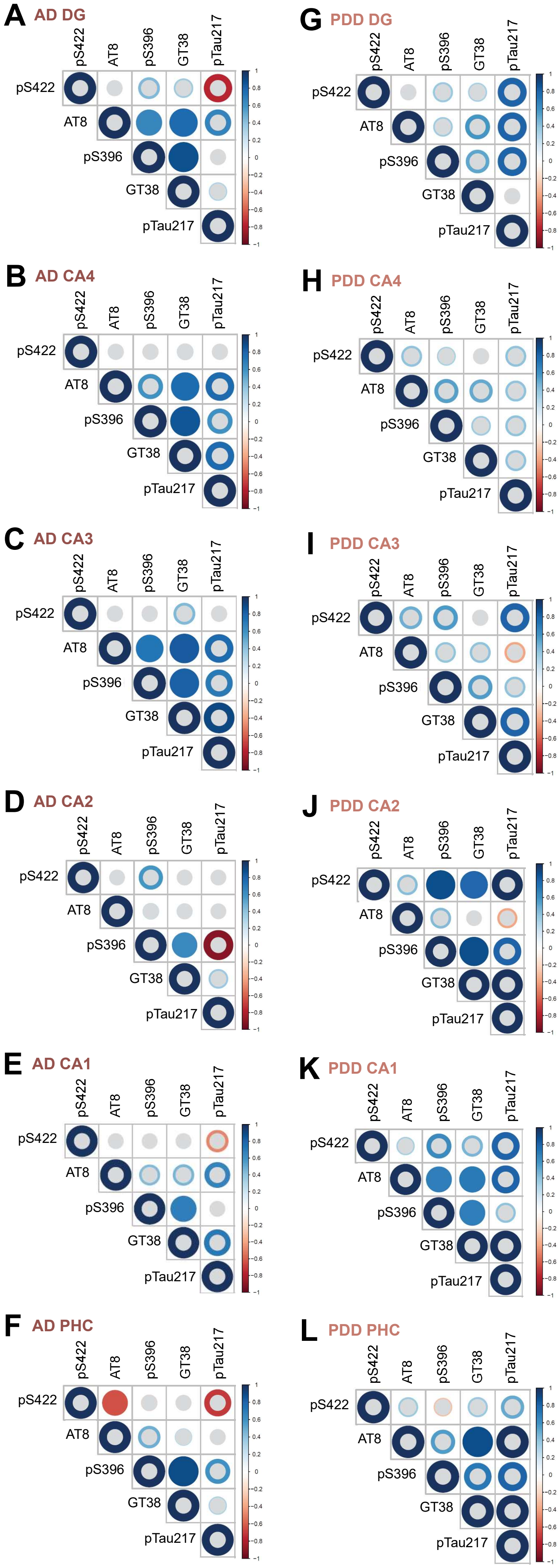


**Supplementary Figure S7**

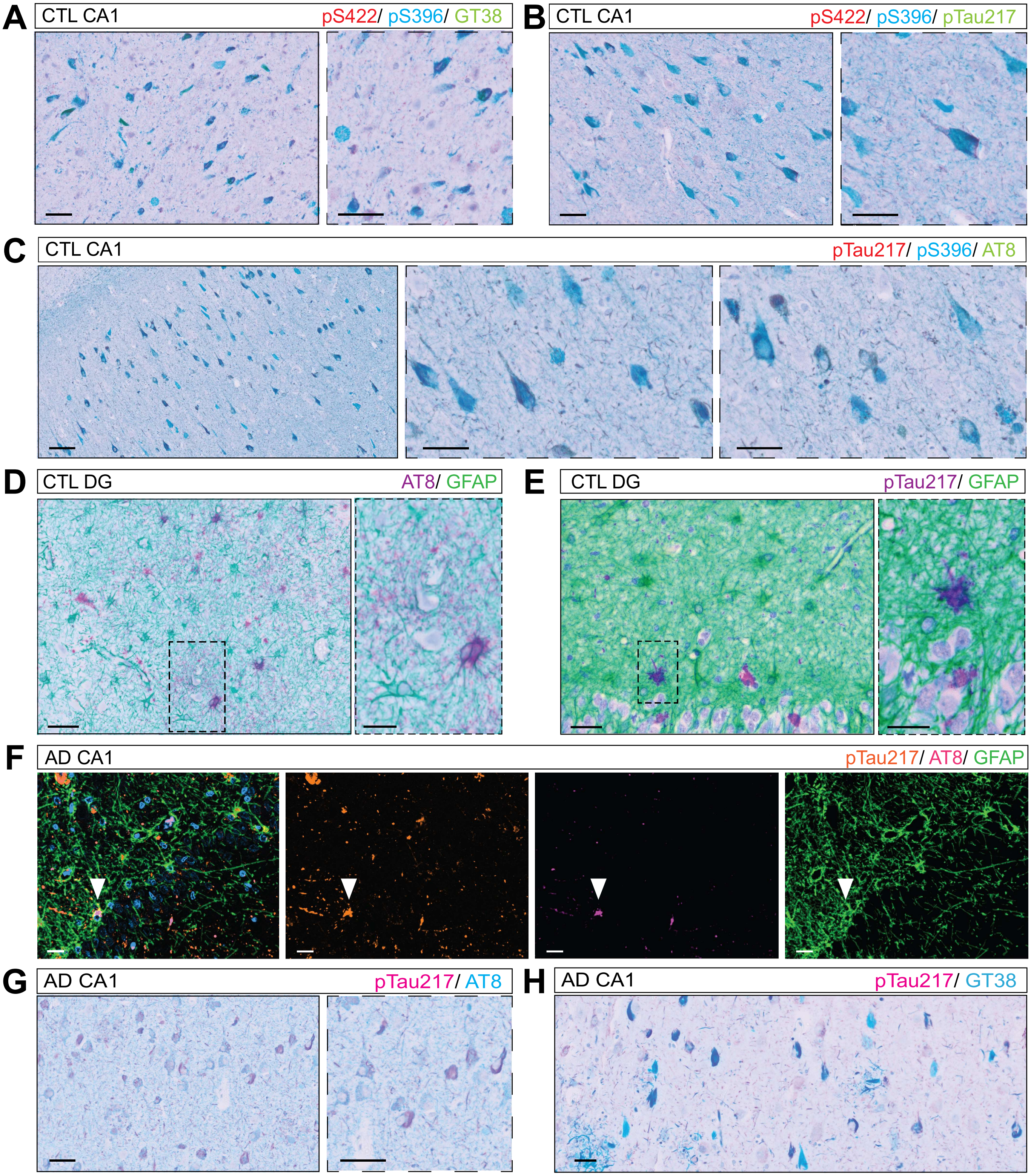


**Supplementary Figure S8**

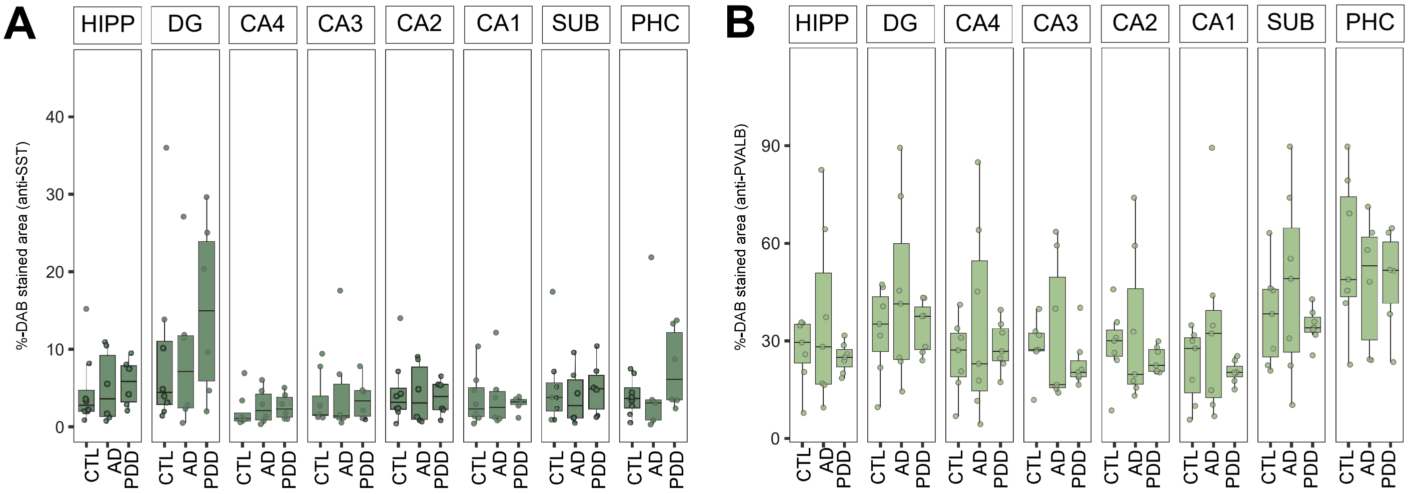


**Supplementary Figure S10**

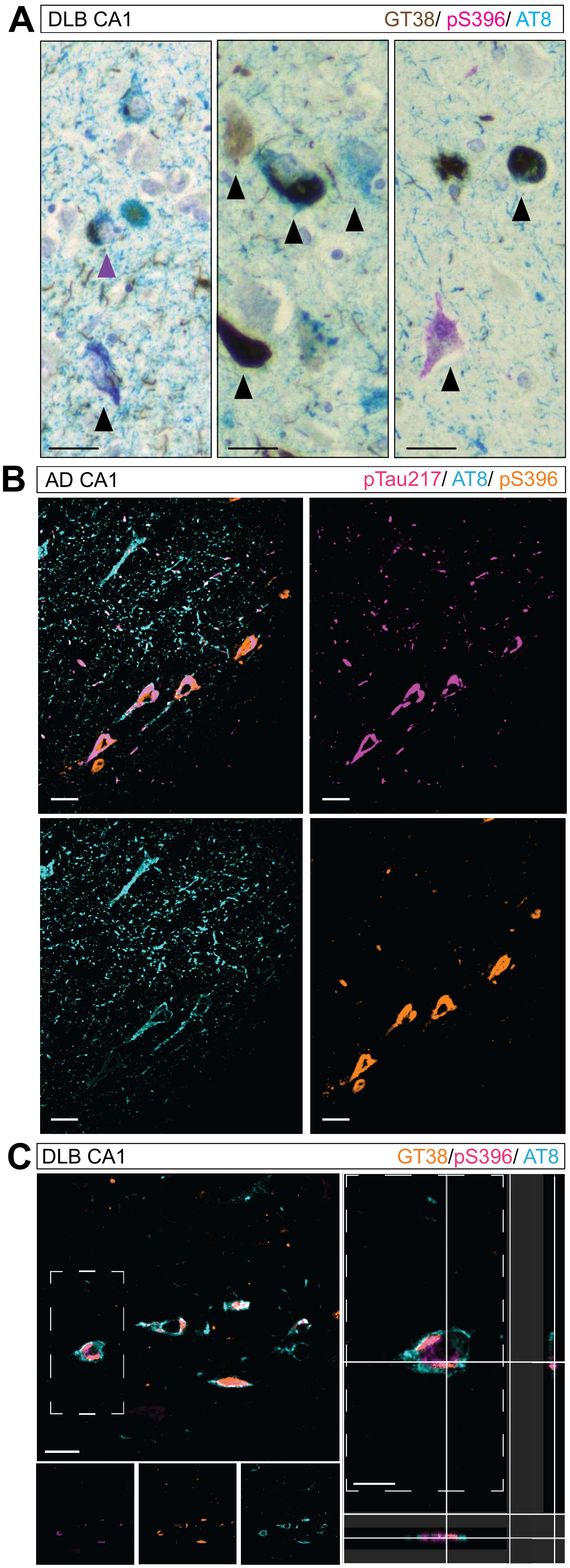


**Supplementary Figure S9**

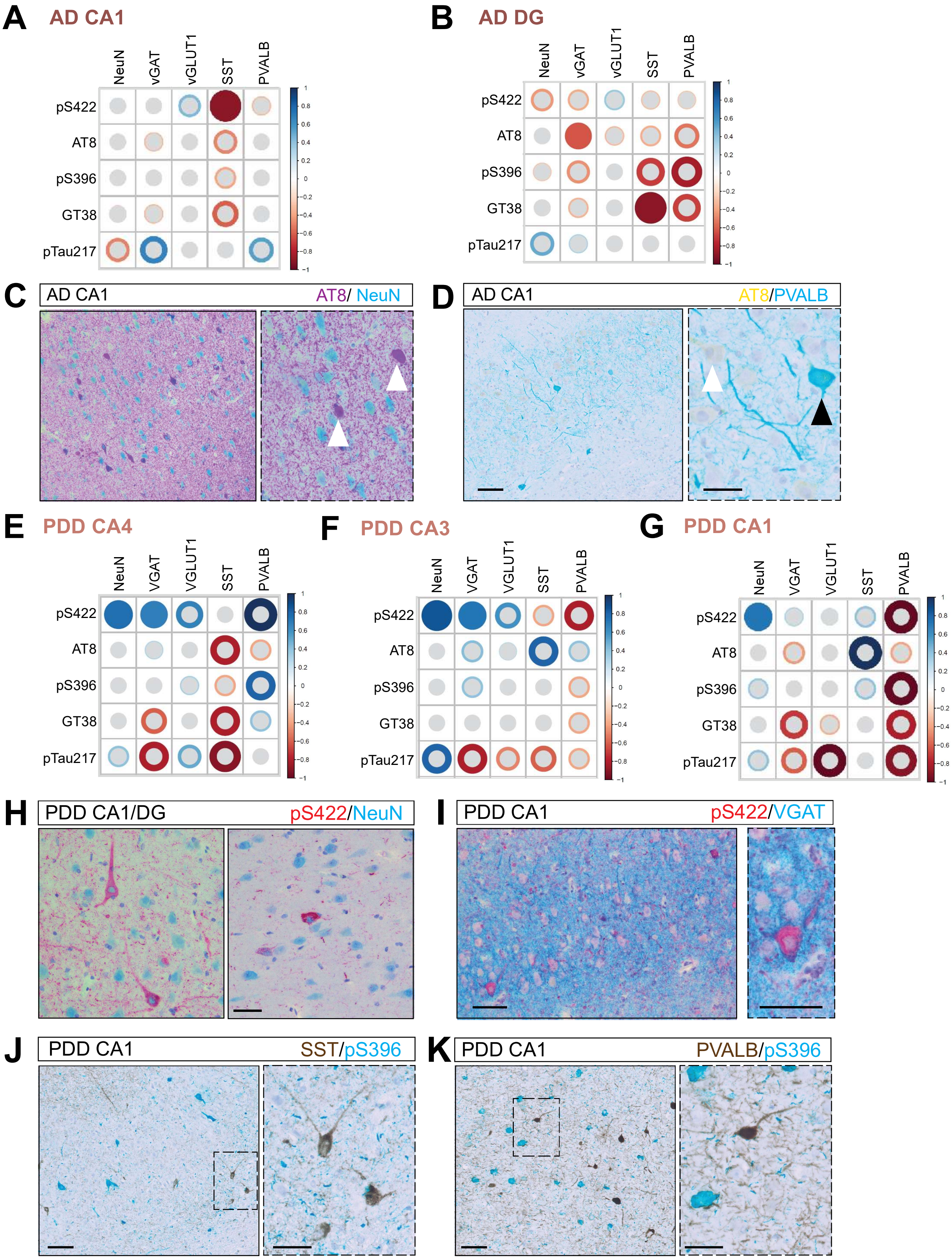


**Supplementary Figure S11**

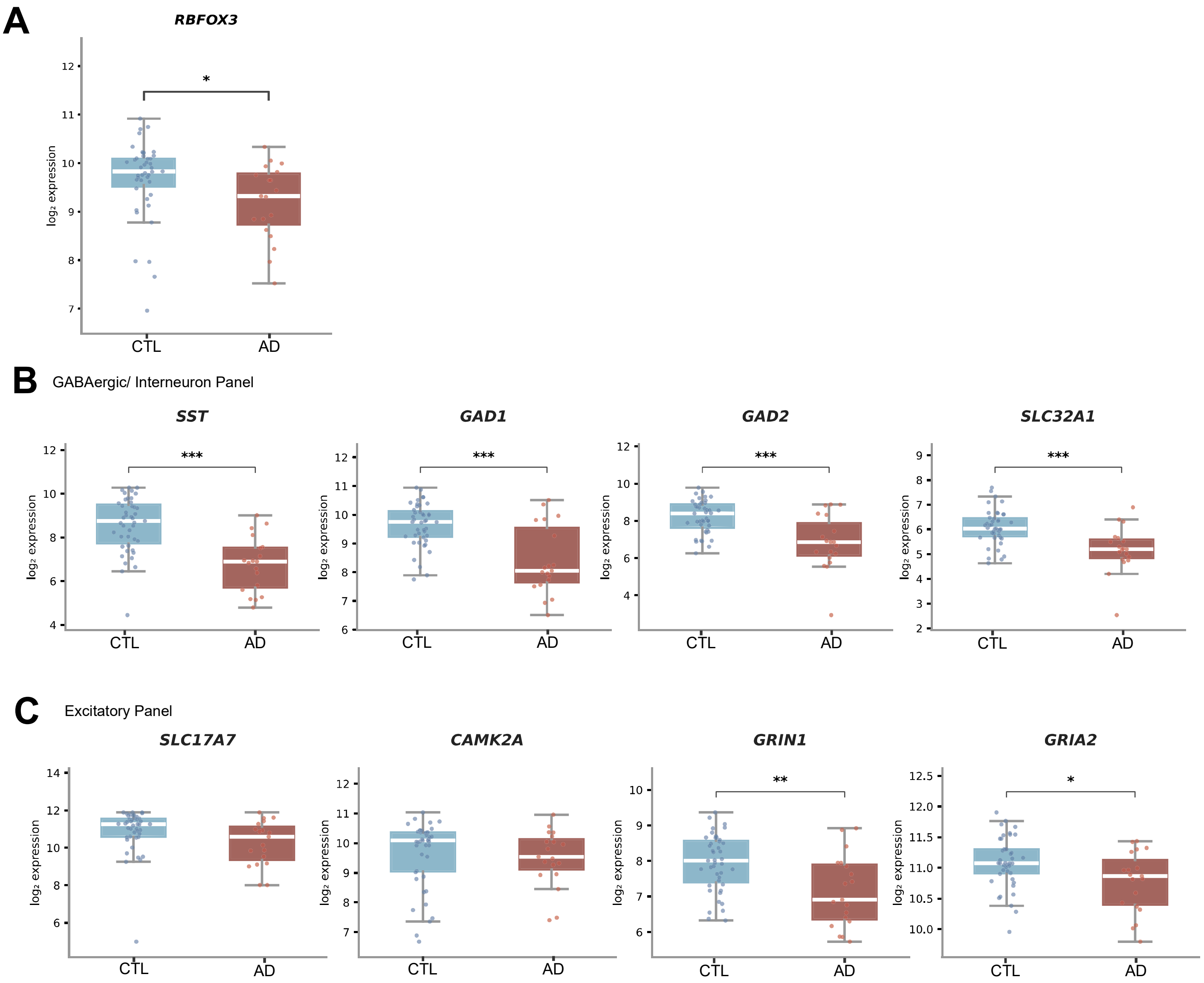


**Supplementary Figure S12**

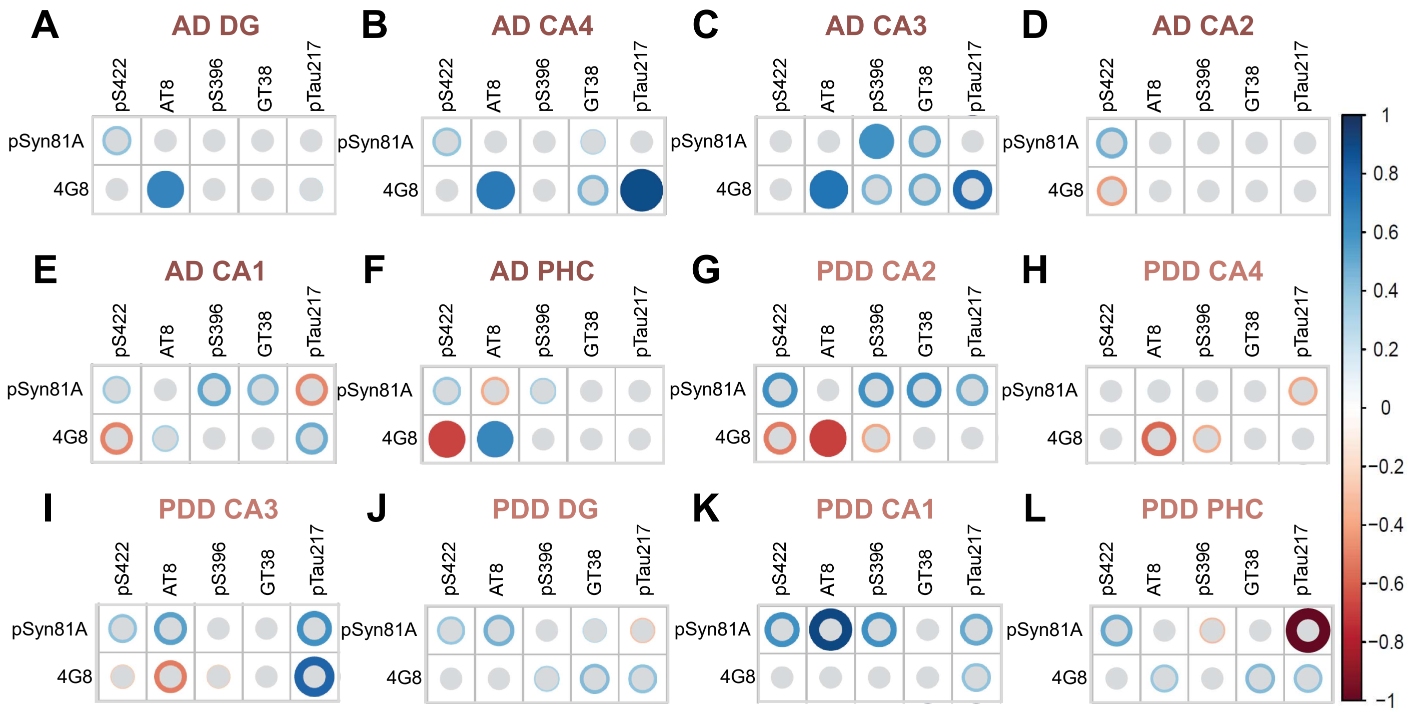


**Supplementary Figure S13**

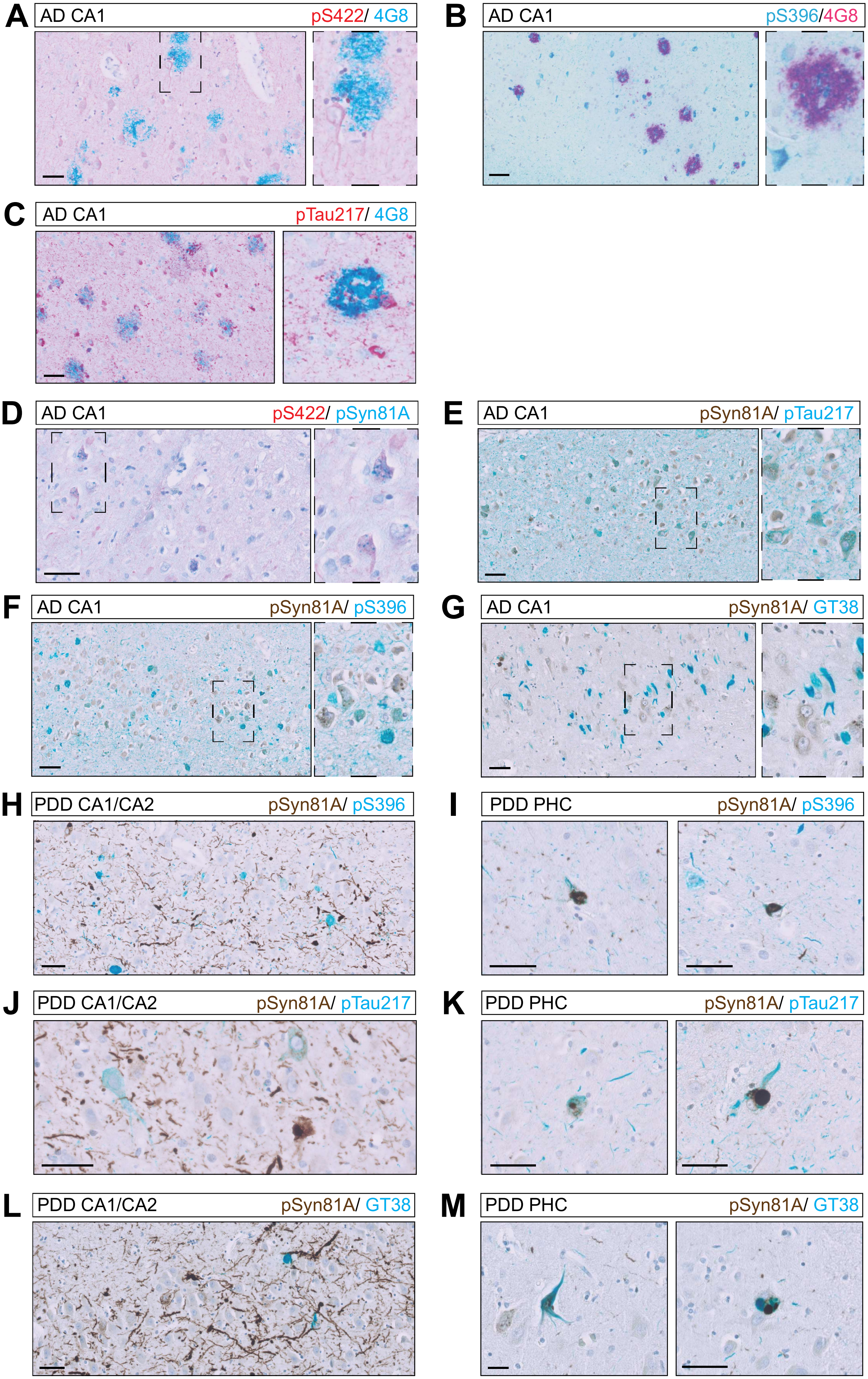


**Supplementary Figure S14**
